# Equivolumetric protocol generates library sizes proportional to total microbial load in next-generation sequencing

**DOI:** 10.1101/2020.02.03.932301

**Authors:** Giuliano Netto Flores Cruz, Ana Paula Christoff, Luiz Felipe Valter de Oliveira

## Abstract

Next-generation sequencing (NGS) has been extensively employed to perform microbiome characterization worldwide. As a culture-independent methodology, it has allowed high-level profiling of sample microbial composition. However, most studies are limited to information regarding relative bacterial abundances, ignoring scenarios in which sample microbe biomass can vary widely. Here, we develop an equivolumetric protocol for amplicon library preparation capable of generating NGS data responsive to input DNA, recovering proportionality between observed read counts and absolute bacterial abundances. Under specified conditions, we argue that the estimation of colony-forming units (CFU), the most common unit of bacterial abundance in classical microbiology, is challenged mostly by resolution and taxon-to-taxon variation. We propose Bayesian cumulative probability models to address such issues. Our results indicate that predictive errors vary consistently below one order of magnitude for observed bacteria. We also demonstrate our approach has the potential to generalize to previously unseen bacteria, but predictive performance is hampered by specific taxa of uncommon profile. Finally, it remains clear that NGS data are not inherently restricted to relative information only, and microbiome science can indeed meet the working scales of traditional microbiology.

## Introduction

The application of next-generation sequencing (NGS) methodologies allows large-scale identification of microorganisms, revealing the colonization and dispersion patterns throughout studied sites such as hospitals, indoor or outdoor natural environments ^1–6^. Despite various detailed microbiome characterization studies, most efforts address solely relative bacterial abundances, *i.e.*, do not account for major variations of total microbial load ^7–9^.

Recent studies claim that the total number of reads in NGS-derived samples (library size) is an arbitrary sum, without biological relevance, yielding microbiome data as necessarily compositional in nature ^8,10,11^. Nonetheless, a previous study has demonstrated that library sizes need not be arbitrary, potentially holding significant correlations with input bacterial cell counts^12^.

The possibility of estimating absolute microbial abundance from NGS data has major impacts for research, government agencies, and industry, empowering researchers and policy makers to address microbiological issues in common scales such as colony-forming units (CFU) without giving up the advantages of high-throughput technology. Further, relative information alone limits decision-making in scenarios in which sample microbe biomass is known to vary widely ^13–15^. Bacterial percentages within a sample are hardly informative in terms surface contamination levels or even risk of microbial environmental dispersion.

The approach herein described was primarily designed for the analysis of samples from indoor environments with varying total biomass (throughout this manuscript we refer to “biomass” as the sample bacterial biomass). Our method enables the improvement of hospital microbiome surveillance as well as other similar sampling sites. Potentially, it can be adapted to broader applications such as clinical evaluations and food safety management. Fixing volumes rather than concentrations during library preparation allows the detection of major variations in input DNA. Under our method, we show that estimation of absolute abundances remains challenged by resolution and taxon-to-taxon variation. We propose Bayesian cumulative probability models to address such issues and demonstrate that total microbial load as well as absolute abundances of observed bacteria can be reliably estimated in terms of CFU.

## Results and Discussion

### Equivolumetric protocol for amplicon library preparation

First, we developed a customized laboratory protocol for amplicon library preparation to recover microbial absolute abundances after NGS sequencing (Figure 1). Briefly, we adapt traditional methods to handle unnormalized inputs of DNA and amplicon. While equimolar protocols standardize samples and PCRs to fixed concentrations ^3,16–20^, we sample equal DNA or amplicon volumes into each library preparation to keep major concentration differences intact. The PCR steps are also optimized for the same purpose, minimizing amplification cycles and stopping before most reactions plateau. Using fewer PCR steps, we decrease error rates and chimera formation, as previously reported ^18,21^. The agarose gel check after library amplifications can still be useful for samples with sufficient biomass, though it is often the case that low biomass samples show no visible bands, hampering any useful interpretation ^12^. For this reason, we do not check for amplicons in agarose gels. In our protocol, PCR pooling is also performed in an equivolumetric fashion, and DNA sequencing follows as in traditional methods. The main justification for our proposal is the fact that samples from indoor environments vary widely in terms of total biomass, generally characterizing low biomass samples ^12–15^ and thus rendering relative information less useful.

**Figure 1.**
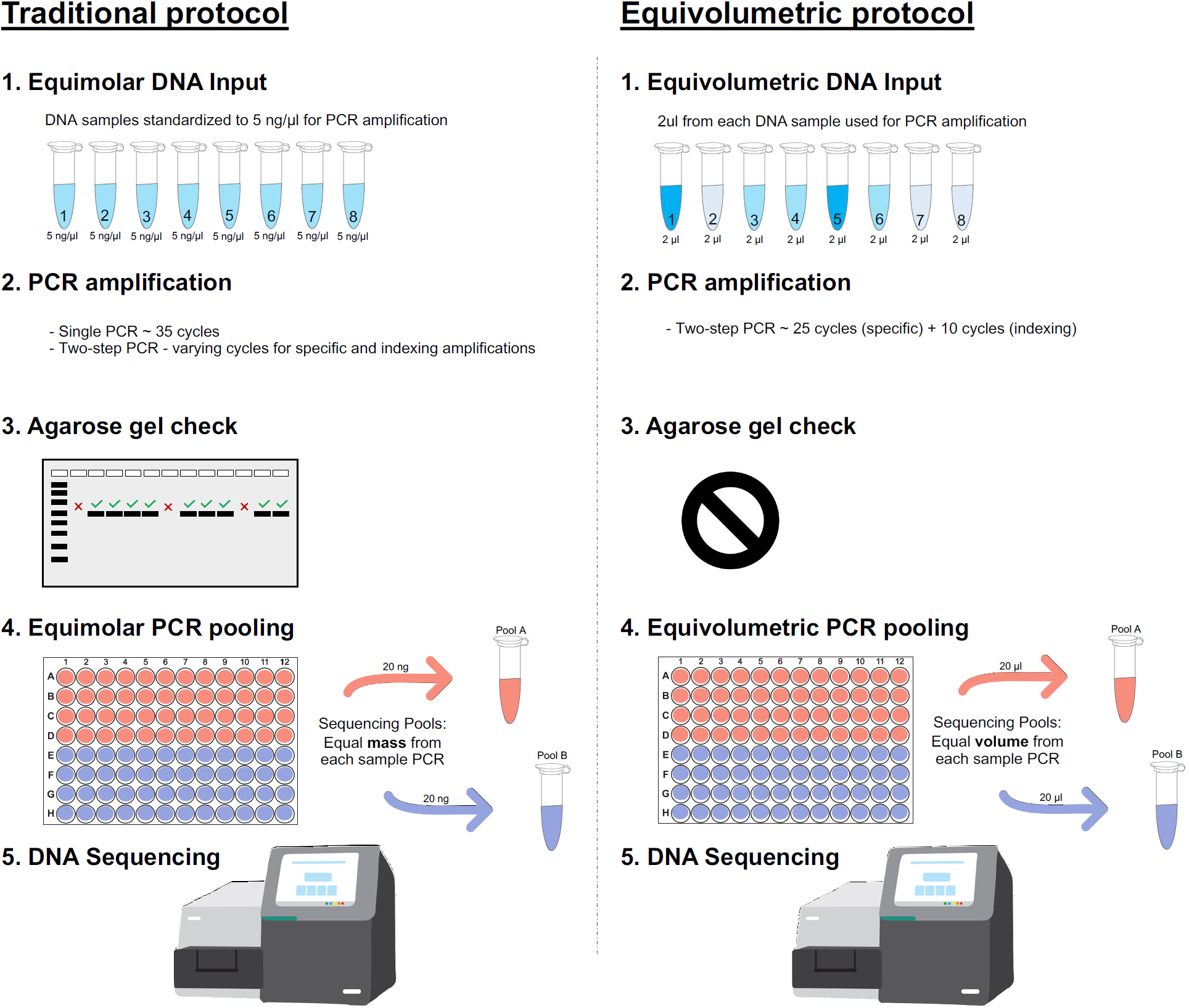
Amplicon library preparation methods for NGS sequencing. **Traditional protocol** is represented as the most common equimolar process. **(1)** Equimolar DNA inputs are prepared based on fluorimetric or spectrophotometric measures, all DNA samples are normalized to equivalent amounts (*e.g*. 5 ng/μL); **(2)** PCR amplifications are performed with single or two-step protocols with varying amplification cycles (most commonly 35 cycles); **(3)** Usually, PCR amplifications are then checked on agarose gel to confirm positive samples and discard negative ones; **(4)** PCR pooling for NGS sequencing is also performed in an equimolar manner through fluorometric quantification (*e.g*. pooling 20 ng from each sample). **Equivolumetric protocol** stands for equal volumes processed for each sample instead of equal concentration. In this protocol, samples retain their original differences in terms of concentrations of input DNA. **(1)** Equal volumes of each sample is used for PCR steps, regardless of its concentration (*e.g*. 2 μL); **(2)** Amplicon library preparation is carried out in a standardized, two-step PCR for 25 cycles using specific marker genes, then additional 10 cycles to add the sequencing adapter and indexes; **(3)** No agarose gel check is performed for these samples since we assume a wide variation in amplicon yield, related to the sample original DNA input; **(4)** PCR pooling for NGS sequencing is performed without specific sample normalizations. Equal volumes are used for each amplicon sample to assemble the NGS sequencing pool (*e.g*. pooling 20 μL from each sample).

### Equivolumetric protocol, input DNA, and absolute bacterial abundances

To investigate whether our approach is capable of recovering absolute abundance information, we first assessed the relationship between NGS generated reads and corresponding input DNA. We used a synthetic DNA molecule with known concentrations (Figure 2A) and sequenced replicated serial dilutions. A polynomial fit demonstrates the sigmoid trend, which indicates NGS-based quantification in absolute terms may still be bounded above by methodological constraints under our protocol (*e.g.*, amplification plateau for highly concentrated samples). We also estimated the corresponding copy numbers and observed similar behavior (Figure 2B). Nonetheless, it is clear from this result that the total number of reads increases with input DNA, which agrees with previous results that showed close relationship between NGS reads and total bacterial cells ^12^.

**Figure 2.**
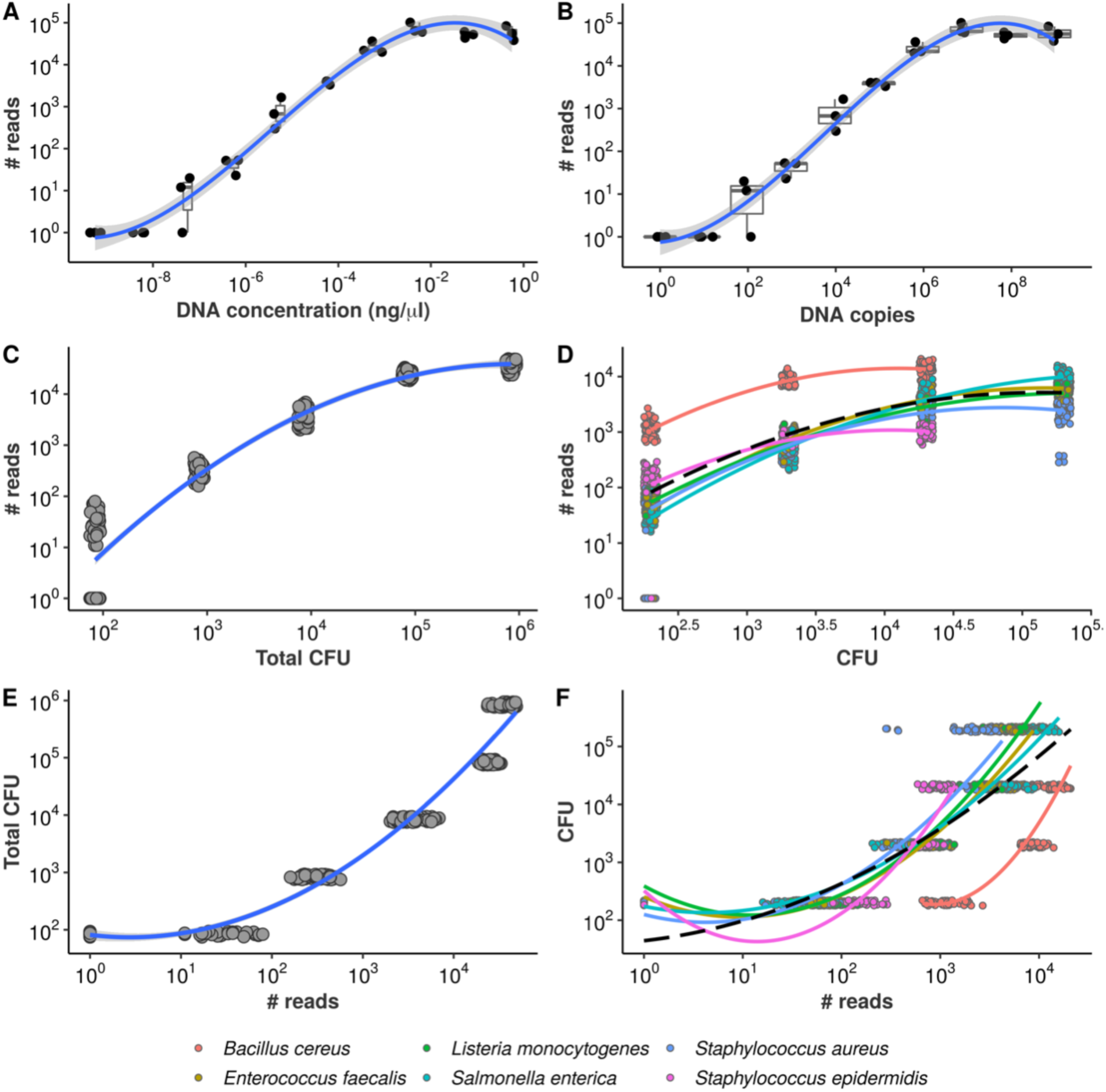
Equivolumetric protocol recovers proportionality between input DNA and NGS reads. Synthetic DNA fragment serially diluted from 0.56 ng/μL to 0.00000056 ng/μL **(A)** or from 954,000,000 to 954 DNA copies **(B)** and its total number of reads obtained by NGS sequencing using the equivolumetric protocol. Total sample reads (library size) from sequencing of serially-diluted samples of mock microbial community using the equivolumetric protocol demonstrates that the obtained read counts are proportional to total microbial load **(C)**. Similar relationship is observed between taxon-specific counts and abundances **(D).** The estimation task of CFU based on NGS reads is illustrated for both total microbial load **(E)** and taxon-specific abundances **(F)**. Total microbial load ranged from 0.84*10^2^ to 0.84*10^6^ CFU, while taxon abundances ranged from 2*10^2^ to 2*10^5^ CFU. A pseudocount of 1 was added to the read counts to avoid *log*_10_(0).

To further confirm the association between reads and sample bacterial load, we performed NGS sequencing in serially-diluted samples of known bacterial concentrations (in terms of CFU), mimicking a surface sample collection from an indoor environment. Our results indicate that the equivolumetric protocol does recover proportionality between NGS data and total microbial load, in terms of both total sample reads (library sizes) (Figure 2C) and bacteria-specific counts (Figure 2D). Hence, our results suggest that compositional constraints motivated by supposedly arbitrary library sizes are not an inherent feature of microbiome datasets, which can indeed be generated to retain absolute abundance information.

The replicates in Figures 2C-D come from four different sequencing runs, demonstrating the reproducibility of the method. As we keep total biomass differences, during data analysis we only correct for variations in the number of reads made available a priori for sequencing in each pool (expected sample coverage, see Methods). Supplementary Material 1 describes data normalization in detail to allow the use of data from multiple sequencing runs as well as an application using real hospital microbiome samples replicated through 14 sequencing runs. We further demonstrate that this approach does not depend on DNA extraction method by testing four different methods with equivalent results (Supplementary Figure 1). While more sophisticated normalizations may be needed for other situations, when varying biomass by orders of magnitudes such step is largely simplified under our protocol.

In practice, microbiome samples are often highly variable in terms of total microbial load. Faecal samples are generally characterized by high biomass, while hospital and indoor samples usually present low biomass ^13,15,22^. Low biomass samples impose more challenges to their processing because of contamination and process inefficiencies ^12^. In fact, in this study we removed *E. coli* sequences from our results since these were frequently detected in our negative controls. We were able to track the corresponding sequences to the DNA polymerase reagent. In fact, *E. coli* was already found to be a common molecular biology contaminant from recombinant enzymes such as polymerases ^23^. Low biomass samples should always be processed especially carefully, accompanied by negative controls to assess possible contaminations ^24–26^.

### Modeling absolute abundance using NGS data

One way to recover absolute bacterial abundance is to associate relative information from high-throughput technology with absolute information from other methods. This has been previously done using qPCR or flow cytometry data as absolute abundance methodologies, and NGS as provider of bacterial proportions ^7,9,27^. Here, however, total microbial load gains importance as a measure of total contamination for surveillance of indoor environments, as opposed to merely a means to project proportions onto absolute terms. In the previous section, we presented data showing NGS reads respond monotonically to the increase of microbial load. We now turn the axes around to describe the present estimation task: given an observed library size (sequenced reads), can total microbial load be predicted reliably? Figure 2E illustrates the problem.

Notice each value of microbial load varies only in orders of magnitude but corresponds to a relatively wide range of observed library sizes. Again, a polynomial trend is fitted, demonstrating the inadequacy of standard linear regression in this case. A similar behavior is observed if we analyze the read counts for each bacterium individually, despite significant taxon-to-taxon variation (Figure 2F). Such naive linear models ignore the monotonic, stepwise fashion in which total microbial load and absolute bacterial abundances vary conditionally on observed reads. Also, there are no prediction bounds: extrapolation towards higher library sizes yields continuously higher predicted values. This is likely unrealistic, given the plateaus observed in Figures 2A and 2B – and the inevitable PCR saturation as total sample biomass increases.

These characteristics led us to consider a cumulative probability model to robustly estimate total microbial load and absolute bacterial abundances using NGS data. In the next subsection, we briefly describe the fitted model for total microbial load – see Methods for formal model specification. We then extend it onto a hierarchical structure that allows variation across bacteria in order to predict taxon-specific absolute abundances. Supplementary Materials 2 and 3 describe both modeling strategies in further detail, including extensive prior and posterior predictive checks as well as assessment of modeling assumptions.

### Cumulative probability model predicts total microbial load

Colony-forming units are continuous measures. However, in classical microbiology, the ability to quantify microbial abundance has limited resolution: values differ mainly in orders of magnitude, and it is difficult to state replicable differences within the same magnitude range. Nonetheless, decision-making based on such data relies mostly on logarithmic differences, and specific orders of magnitude are often the main interest – *e.g.* diagnosis of urinary tract infection^28^, bacterial characterization from International Space Station surfaces ^19^, or hospital environments^29^.

In this scenario, we propose a cumulative probability model (CPM) with logit link, also known as Proportional Odds (PO) model, to predict total microbial load in terms of colony-forming units (CFU) based solely on library size. CPMs are capable of robustly estimating discrete and continuous outcomes in a semi-parametric fashion, both within Frequentist and Bayesian frameworks ^30,31^. Another advantage is the wealth of information produced: by modeling the cumulative probability function conditional on the data, one can retrieve predictions in terms of class of highest probability (the most likely outcome, herein referred to as CHP), expected values, tail probabilities, and even quantiles ^32^. See Methods for the entire (Bayesian) model specification and Supplementary Material 2 for the detailed workflow.

We fit the model using the R package brms ^33^, and Figures 3A-3D show the results. Figure 3A shows the estimated class probabilities as a function of library size, and Figure 3B shows the implied expectations (black solid line, 95% credible intervals in blue) as well as the CHP (red solid line). Predictive intervals for the CHP are also shown in light grey. Notice how the red line in Figure 3B follows closely the behavior of the class probabilities in Figure 3A. The predictions generated by the ordinal model are by construction monotonic. Also note that both expectation and CHP are bounded within the observed outcome space, overcoming extrapolation issues related to the previous naive model. In Figure 3C, posterior-predictive check indicates the overall structure of the observed data is well captured by the posterior draws of the model.

**Figure 3.**
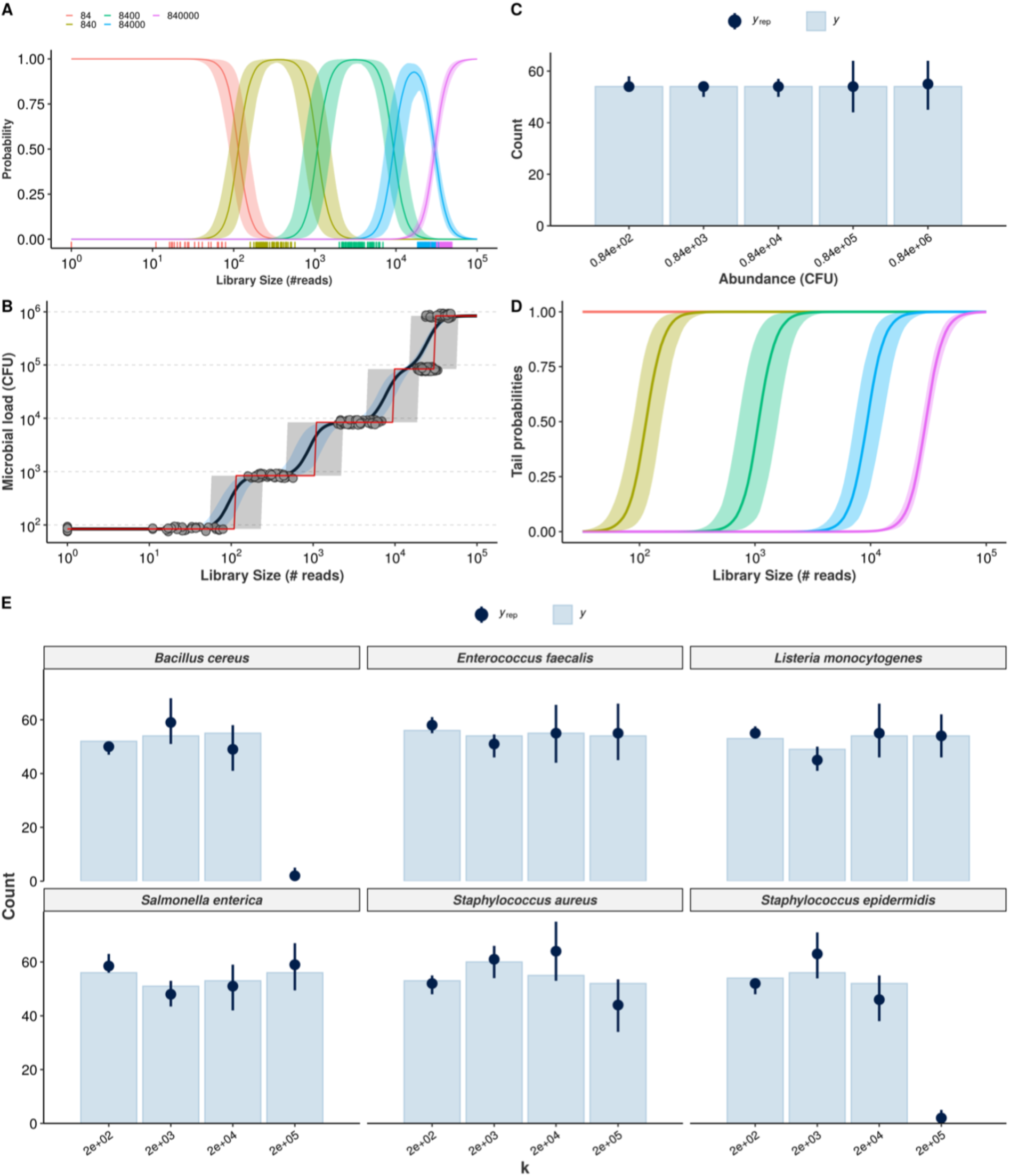
Cumulative probability models for the estimation of absolute bacterial abundances. Estimation of class probabilities for each observed value of total microbial load (in CFU), conditional on observed library size, is retrieved from the ordinal logistic regression framework **(A)**. Conditional expectations are then derived as weighted average of microbial load values and respective class probabilities (black line, 95% credible intervals in light blue) **(B)**. The class of highest probability (CHP, the most likely outcome given the observed reads) is also shown (red line, 95% predictive intervals in grey). Posterior predictive check shows the Bayesian model captures the overall structure of the observed data for total microbial load (*y*_*rep*_: posterior draws, *y*: observed data) **(C)**. Tail probabilities, herein defined as the probability of observing at least class ck, conditional of observed library size are an alternative for cases in which CHP- and expectation-based predictions are prohibitively uncertain **(D)**. Hierarchical CPM accounts for differences across bacteria and takes advantage of partial pooling to estimate taxon-specific abundances **(E)**. The resulting posterior predictive check indicates no major signs of misfit.

Finally, Figure 3D shows model-implied tail probabilities, herein defined as the probability of observing at least abundance *c*_*k*_, *i.e., Pr*(*Y*_*i*_ ≥ *c*_*k*_|*X* = *x*_*i*_) rather than *Pr*(*Y*_*i*_ > *c*_*k*_|*X* = *x*_*i*_). The ability of deriving lower/upper bounds with high probabilities is a major advantage of CPM. Under this model, even though distinguishing abundance values may be challenging for certain ranges of the predictor space, one can still rely on tail probabilities to guide decision-making. For instance, should CHP- and expectation-based predictions be shown limited in performance, or the predictive interval prohibitively wide, then one can still use the highest class *c*_*k*_ such that the probability of having at least*c*_*k*_ CFU is no less than a given probability threshold *τ, i.e*., for each observation *i* find 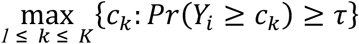 for some large *τ* (*e.g.* 95%).

In the next subsection, we extend the previous model to handle taxonomic information in a hierarchical fashion so that one can make predictions of absolute abundance for each bacterium individually. We then validate both models using cross-validation and prediction on held-out samples (test sets) in the following section.

### Hierarchical CPM predicts absolute bacterial abundances

In order to handle taxonomic information, we formulate a similar model which incorporates taxon-specific effects in a hierarchical structure. The major difference is that the linear predictor term is parametrized with population-level parameters and group-level counterparts (for both intercept and slope), allowing predictions of many bacteria with a single model and taking advantage of partial pooling ^34^.

While we use seemingly weakly-informative priors (see Methods for full model specification), their joint behavior favors outer classes to improve distinguishability when dealing with classes of overlapping average number of reads. This is illustrated with prior-predictive check and assessment of CPM assumptions in Supplemental Material 3, which also shows detailed workflow and visualizations for each observed bacteria. Our prior choice resulted from model comparison with approximate leave-one-out cross-validation ^35,36^. Figure 3E shows the corresponding posterior predictive check, suggesting the data is well captured by the model-implied data generation process for all bacteria. *B. cereus, S. aureus*, and *S. epidermidis* may represent challenging cases, although this can be an artifact from estimating varying effects with only six bacteria (stronger priors led to worse fit during model comparison).

### Model validation

We validated both models using 10-fold cross-validation (CV) and prediction on held-out test sets comprising of 10% of the total number of observations. We assess performance both as classification and regression tasks, using CHP- and expectation-based predictions.

Figure 4A shows the 10-fold CV results for the total microbial load model. For visualization, we have split the assessed metrics into bounded between 0 and 1 and unbounded metrics. Bounded metrics based on CHP included the observed coverage of 95% predictive interval, Somers’ Delta (measure of ordinal association), classification accuracy, and Spearman’s rank correlation. The latter was also assessed for expectation-based predictions. In general, these metrics varied well above 0.9. Notably, the predictive intervals showed 100% coverage, which is likely overconfident. Nonetheless, most intervals spanned two abundance classes as in Figure 3B (see also Supplementary Materials 2 and 3), suggesting errors occur mainly within one order of magnitude from the true values. Ordinal association, as measured by Somers’ Delta, was consistently greater than 0.95 for both CV and test set.

**Figure 4.**
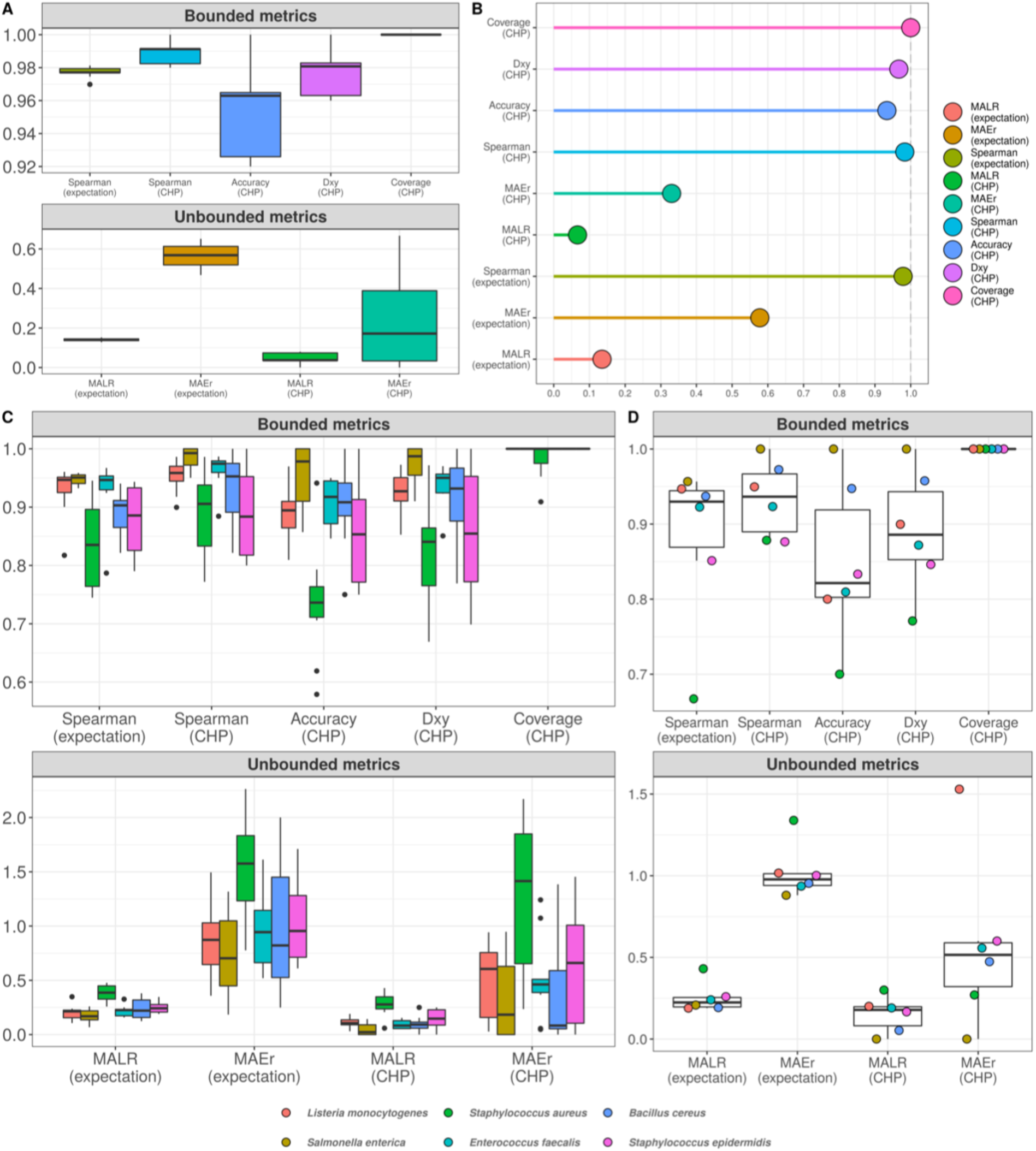
Cumulative probability models generate accurate predictions for total microbial load and taxon-specific absolute abundances. Performance measures from 10-fold cross-validation of total microbial load model indicate predictive errors are constrained far below one order of magnitude **(A**). For visualization, bounded metrics vary between 0 and 1, while unbounded metrics vary in the positive real line. Similar results were observed in the held-out test set **(B**). 10-fold cross validation for taxon-specific predictions using hierarchical CPM indicates predictive performance varies across bacteria, although still far below one order of magnitude **(C**). Similar results were observed in the held-out test set **(D)**. Predictions based on class of highest probability are indicated with (CHP) in the x axis, and expectation-based counterparts are indicated likewise. MALR: mean absolute log-ratio; MAEr: mean absolute error relative to true values; *D*_*xy*_: Somers’ Delta measure of ordinal association; Coverage: observed coverage of 95% predictive interval.

Unbounded metrics relied on modified versions of absolute errors, for both CHP- and expectation-based predictions. MALR and MAEr denote Mean Absolute Log-Ratio and Mean Absolute Error relative to true value, respectively, defined as follows.

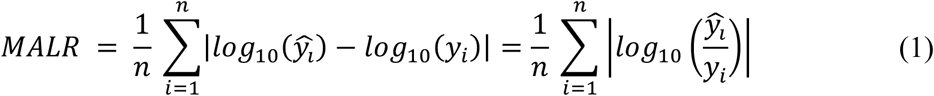

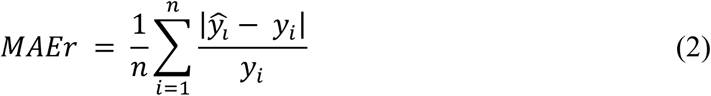

MALR represents deviance in orders of magnitude, which varied during CV below 0.2 for both CHP and expectation, a result reproduced in the test set evaluation (Figure 4B). Perhaps more intuitive, MAEr tended to be smaller for CHP-based predictions compared to expectations. During 10-fold CV or test-set validation, we did not observe MAEr values greater than 0.7.

Although not as common as mean absolute errors or mean squared errors, the metrics herein assessed do not penalize estimation in varying orders of magnitude, offering advantages in interpretation. A MALR value of 1 corresponds to a ratio between predicted and observed values of one order of magnitude in the *log*_*10*_ scale. A MAEr of 1 indicates prediction absolute error as large as the true value, which would still be largely insignificant given the logarithmic scale. Using both CHP- and expectation-based predictions, our results indicate that predictions for the total microbial load model were mostly kept within the observed orders of magnitude.

Figures 4C (10-fold CV) and 4D (test-set) show the analogous measures for the hierarchical model with bacteria-specific predictions. Median MAEr varied below 1 for both CHP and expectations during CV for most bacteria. In the test set, the highest value observed was 1.5 (*L. monocytogenes*). We observed most median accuracy values as high as 0.85 during CV and 0.8 for the test set, while ordinal association seems slightly higher in general. The model was least performant for predicting *S. aureus* abundance, as indicated by almost all metrics computed. Still, observed MALR varied consistently below 0.5 for all bacteria both in CV and test-set validation. Again, the results indicate our predictions are contained within respective orders of magnitude, suggesting that NGS reads can indeed be a valuable source of information regarding absolute bacterial abundances.

### Predicting abundance of previously unseen bacteria

As the hierarchical CPM enables prediction of previously unseen bacteria, we also performed leave-one-group-out CV to assess how our model could generalize in high-throughput settings, in which one may have dozens of taxa of interest. For each bacterium, we hold out its corresponding data points and train a separate model with the remaining data. We then perform predictions for the held-out taxon, treating it as “previously unseen” - not used during model fitting.

Figure 5 shows the results. Classification accuracy drops substantially, and the model completely fails to classify abundance values for *B. cereus*. Yet, for other bacteria, ordinal association and classification accuracy varied between 0.9 and 0.6. Treated as a classification task, the poor predictive performance is likely influenced by high uncertainty associated with the estimation of varying intercepts and slopes using data from only five bacteria at each iteration. This can also explain the high coverage values: predictive intervals were so wide that potentially spanned nearly all outcome space. On the other hand, the taxon associated with the worst out-of-sample performance (*B. cereus*) was also the one with the highest random effects in the original model, *i.e.*, the greatest deviance from the overall, population-level effects – see Supplementary Material 3. While most bacteria showed MAEr between 0.5 and 3, *B. cereus* exceeded the value of 50 (absolute error as large as 50 times the true value). Nonetheless, MALR still varied below the threshold of 1 for all but *Bacillus cereus*, which almost reached a MALR of 2 (two orders of magnitude).

**Figure 5.**
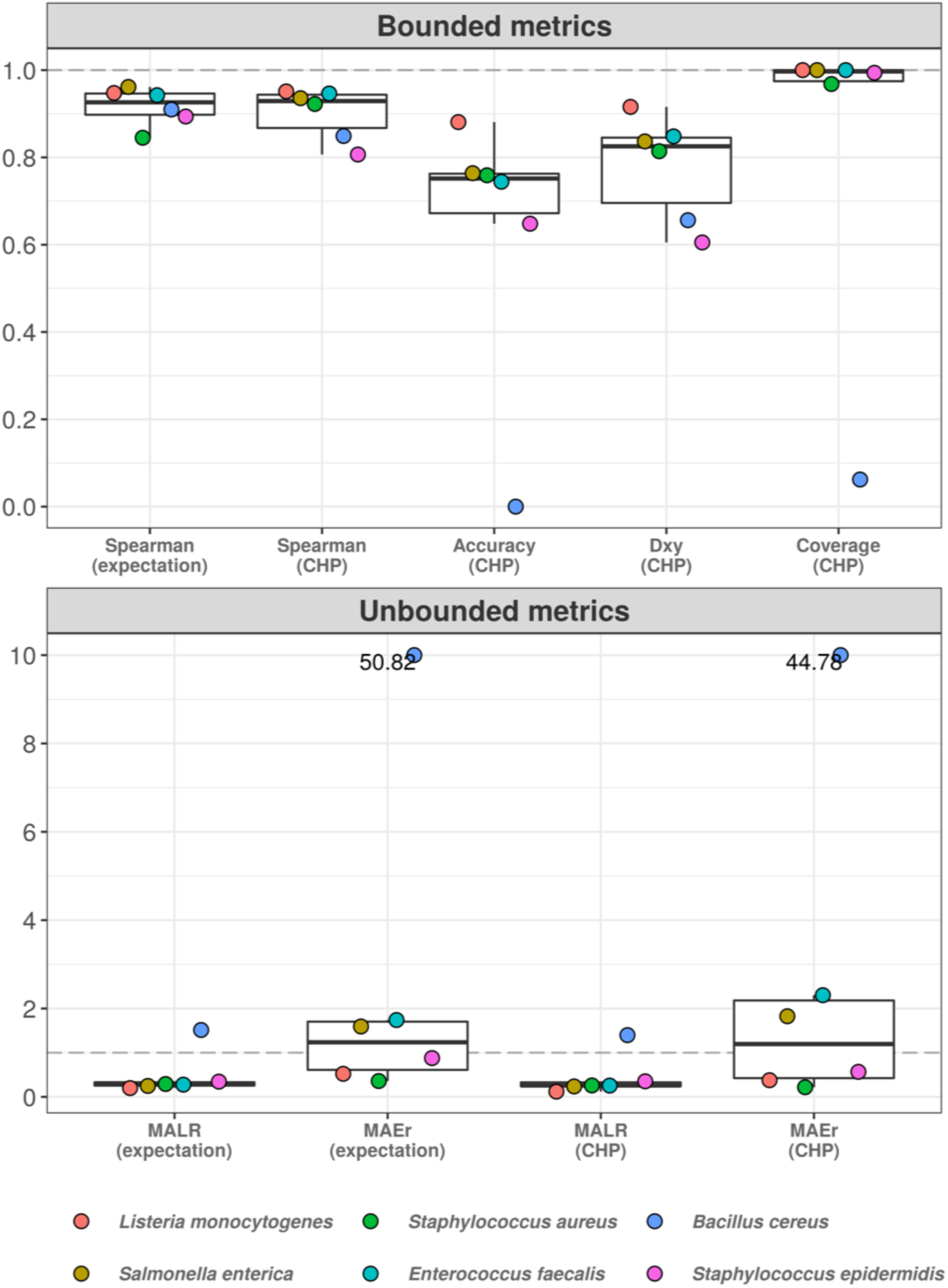
Hierarchical cumulative probability model predicts previously unseen bacteria with varying performance. Leave-one-group-out cross-validation was used to estimate predictive performance of hierarchical CPM for previously unseen bacteria. The predictive errors are constrained below one order of magnitude for most bacteria, except for *Bacillus cereus* – which reached errors of almost two orders of magnitude (lower panel). The dashed grey line indicates a value of 1, representing one order of magnitude in the context of MALR. The model fails to classify abundance values of *Bacillus cereus* (upper panel), although ordinal association (*D*_*xy*_) remains above 0.6. Most absolute errors represent no more than two times the observed abundances in a context of logarithmic differences – except for *B. cereus* and (slightly) *E. faecalis* using CHP. For visualization, we truncated the y axis of the lower panel at the value of 10 and indicated higher values with numeric labels.

Thus, it is clear that generalization in high-throughput settings is challenged by specific bacteria, such as *B. cereus*, which deviate greatly from overall profiles. Yet, prediction errors for most bacteria were shown to vary below one order of magnitude, suggesting the approach is promising. Improvements can also be achieved, for instance, through the inclusion of more predictors such as 16S rRNA gene copy number, gram-like classifications, and even taxonomic information from higher ranks. The addition of more bacteria species to estimate varying effects can also be beneficial.

Overall, our results indicate the predictive errors for CFU do not exceed one order of magnitude (on the *log*_*10*_ scale) for observed bacteria. While total microbial load seems more reliably estimated, for both models the absolute errors tend to be no greater than two times the true values – in a reality of logarithmic differences. While one might doubt the importance of estimating *4* * *10*^*3*^ CFU compared to a true value of *2* * *10*^*3*^ CFU, we acknowledge our models can be improved by adding more data points, predictors, and different bacteria. Yet, it remains clear that library sizes need not to be arbitrary, and that NGS reads can indeed produce reliable estimations of absolute bacterial abundances, at least in the working scales of classical microbiology.

## Conclusion

Here we have shown that assessment of absolute bacterial abundance using NGS data becomes possible upon protocol alteration within specified conditions, and the remaining challenges lie within the realm of resolution and taxon-to-taxon variation. While library sizes do depend on sequencing effort – a partially arbitrary sequencing setup -, equivolumetric protocols assure the maintenance of major input DNA variations, at least for certain ranges of colony-forming units. Under such conditions and assuming the same protocol for a set of samples of similar nature, our results indicate that these procedures recover the proportionality between library sizes and total microbial load.

We have also developed Bayesian cumulative probability models to robustly estimate both total microbial load and bacterial absolute abundances using NGS data only. Most prediction errors lied far below the threshold of one order of magnitude, indicating that the models are sufficiently reliable. Still, further research is needed to understand whether such models can generalize to high-throughput settings, in which data from a small subset of taxa are used to make predictions on previously unseen bacteria. Finally, it is clear that the claim that library size is always an arbitrary sum, often taken for granted by several previous works, is readily overcome by the methods herein proposed.

## Methods

### Samples

A synthetic DNA fragment with a naturally non-occurring sequence was designed with the 16S rRNA V3/V4 primers sequences flanking their extremities. This fragment with 544 bp was synthesized as gBlocks® Gene Fragments from IDT (IA, USA). This DNA was eluted to a final concentration of 10 ng/μL in TE buffer following the manufacturer instructions. Then it was serially diluted from 0.56 ng/μL to 0.00000056 ng/μL by a 10X factor dilution. This serial dilution is equivalent to a range of 954,000,000 to 954 copies of the synthetic DNA. Samples were processed in experimental triplicates.

Reference bacterial isolates were acquired from ATCC (American Type Culture Collection, VA, USA) ATCC 19111 *Listeria monocytogenes*, ATCC 14028 *Salmonella enterica*, ATCC 10876 *Bacillus cereus*, ATCC 12228 *Staphylococcus epidermidis*, ATCC 29212 *Enterococcus faecalis*, ATCC 8739 *Escherichia coli*, ATCC 25923 *Staphylococcus aureus*. These bacterial isolates were individually grown overnight at 35°C in Brain Heart Infusion media and then adjusted to an optical density (OD_600_) of 0.5 to be further diluted with a 10X factor for more seven consecutive dilutions. The two more diluted concentrations for each bacterium had 100 μL plated in PCA (Plate Count Agar) and incubated overnight at 35°C to check for the CFU (Colony Forming Units) concentrations used in the assay described below. The dilutions corresponding to 2, 20, 200, 2000, 20000 and 200000 CFU for each above bacteria were inoculated in a sterile plastic plate, without media, and left to dry in a biological safety cabinet. Then the bacterial cells were collected from the dry plate surface using a sterile hydraflock swab (Puritan, ME, USA) moistened with sterile physiological solution. After sample collection the swab was broken down into a microtube containing 800 μL of stabilization solution – ZSample (BiomeHub, SC, BR). The DNA from the above collected samples was extracted using a thermal lysis protocol along with AMPure XP magnetic beads purification (Beckman Coulter, CA, USA). Samples were processed with fifteen replicates for each bacterial CFU dilution. Additionally, three alternative DNA extraction kits were used: QIAamp® DNA Mini and Blood Mini (QIAGEN, Germany), lot: 154018620, DNAeasy Power Soil (QIAGEN, Germany), lot: 163024722 and DNAeasy Power Soil PRO (QIAGEN, Germany), lot: 160048809.

### Library preparation and sequencing

The 16S rRNA amplicon sequencing libraries were prepared using the V3/V4 primers (341F CCTACGGGRSGCAGCAG and 806R GGACTACHVGGGTWTCTAAT) ^37,38^ in a two-step PCR protocol. The first PCR was performed with V3/V4 universal primers containing a partial Illumina adaptor, based on TruSeq structure adapter (Illumina, USA) that allows a second PCR with the indexing sequences similar to procedures described previously ^17^. Here, we add unique dual-indexes per sample in the second PCR. Two microliters of individual sample DNA were used as input in the first PCR reaction. The PCR reactions were carried out using Platinum Taq (Invitrogen, USA) with the conditions: 95°C for 5 min, 25 cycles of 95°C for 45s, 55°C for 30s and 72°C for 45s and a final extension of 72°C for 2 min for PCR 1. For PCR 2, two microliters of the first PCR were used and the amplification conditions were 95°C for 5 min, 10 cycles of 95°C for 45s, 66°C for 30s and 72°C for 45s with a final extension of 72°C for 2 min. All PCR reactions were performed in triplicates. The second PCR reactions were cleaned up using AMPureXP beads (Beckman Coulter, USA) and an equivalent volume of each sample was added in the sequencing library pool. At each batch of PCR, a negative reaction control was included (CNR). The final DNA concentration of the library pool was estimated with Quant-iT Picogreen dsDNA assays (Invitrogen, USA), and then diluted for accurate qPCR quantification using KAPA Library Quantification Kit for Illumina platforms (KAPA Biosystems, MA). The sequencing pool was adjusted to a final concentration of 11.5 pM (for V2 kits) or 18 pM (for V3 kits) and sequenced in a MiSeq system (Illumina, USA), using the standard Illumina primers provided by the manufacturer kit. Single-end 300 cycle runs were performed using V2×300, V2×300 Micro, V2×500 or V3×600 sequencing kits (Illumina, USA) with sample coverages specified in Supplementary table 1.

### Bioinformatics analysis and taxonomic assignment

The sequenced reads obtained were processed using the bioinformatics pipeline described below (BiomeHub, SC, BR – hospital_miccrobiome_rrna16s:v0). First, Illumina reads have the amplicon forward primer checked, it should be present at the beginning of the read, and only one mismatch is allowed in the primer sequence. The whole read sequence is discarded if this criterion is not met. The primers are then trimmed, and the reads accumulated error evaluated. Read quality filter (*E*) is performed converting each nucleotide Q score in error probability (e_i_), that is summed and divided by read length (L).

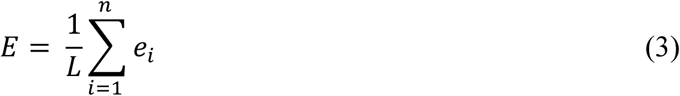

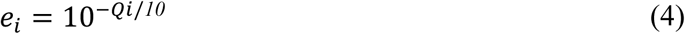

Reads are then analyzed with the Deblur package v.1.1.0 ^39^ to remove possible erroneous reads and identical sequences are grouped into oligotypes (clusters with 100% identity). The sequence clustering with 100% identity provides a higher resolution for the amplicon sequencing variants (ASVs), also called sub-OTUs (sOTUs) ^40^ – herein denoted as oligotypes. Next, VSEARCH 2.13.6 ^41^ are used to remove chimeric amplicons. We implemented an additional filter to remove oligotypes below the frequency cutoff of 0.2% in the final sample counts.

We also implemented a negative control filter for low biomass samples. If any oligotypes are recovered in the negative control results, they are checked against the samples and automatically removed from the results only if their abundance (in number of reads) are no greater than two times their respective counts in the sample. The remaining oligotypes in the samples are used for taxonomic assignment with the BLAST tool ^42^ against a reference genome database (encoderef16s_rev6_190325, BiomeHub, SC, Brazil). This database is constructed with complete and draft bacterial genomes, focused on clinically relevant bacteria, obtained from NCBI. It is composed of 11,750 sequences including 1,843 different bacterial taxonomies.

Taxonomy are assigned to each oligotype using a lowest common ancestor (LCA) algorithm. If more than one reference can be assigned to the same oligotype with equivalent similarity and coverage metrics (e.g. two distinct reference species mapped to oligotype “A” with 100% identity and 100% coverage), the taxonomic assignment algorithm leads the taxonomy to the lowest level of possible unambiguous resolution (genus, family, order, class, phylum or kingdom), according to similarity thresholds previously established ^43^.

### Normalization

The normalization procedure is fully described in Supplementary Material 1. Briefly, let 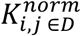 denote the normalized counts for the taxonomy *i* sample *j* ∈ *D*, where *D* is the set of samples from the *d* ^*th*^ sequencing run. Then the normalization is a simply rescaling of the raw counts.

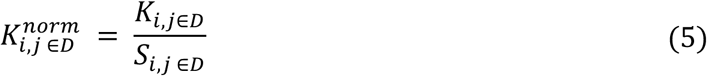

The size factor is sequencing-specific and is calculated as follows:

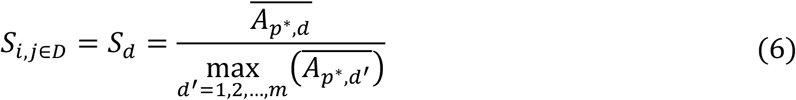

where 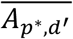 is the average number of reads per sample made available a priori in the sequencing pool of interest *p*^⋆^ within sequencing *d* (expected sample coverage). The lower the relative availability, the smaller the resulting factor and thus greater the normalized values relative to the raw counts. Once normalized, divergences across samples from different sequencing runs, but of similar bacterial abundances, are assumed to rise mostly from sequencing efficiency differences – yet of negligible order of magnitude.

### Statistical Analysis

All statistical analyses were performed using R software environment version 3.6.2 ^44^. We used the brms R package and Stan (v. 2.11.1 and v. 2.19.1, respectively) to perform all bayesian analyses and the tidyverse package suite (v. 1.3.0) for data wrangling and visualization ^33,45,46^. We also used the phyloseq R package (v. 1.30.0) to handle microbiome data ^47^. Supplementary Table 2 lists all R packages used along with corresponding versions and references. The entire modeling strategy is further detailed in Supplementary Material 2 (total microbial load) and 3 (bacterial abundances). All models were fit within the Bayesian framework.

### CPM for total microbial load estimation

We used a cumulative probability model (CPM) with a logit link, also known as Proportional Odds (PO) model, to predict total microbial load based on NGS reads. Let *Y*_*i*_ denote the total microbial load (in CFU scale) from the *i*^*th*^ sample. Given our serially-diluted samples, we only observe *K* = 5 abundance values such that *Y*_*i*_ takes values *c*_*k*_ ∈ {*c*_1,_ *c*_2,_ …, *c*_*K*_} = {0.84 × 10^2,^ 0.84 × 10^3^, …, 0.84 × 10^6^}. We then define the model:

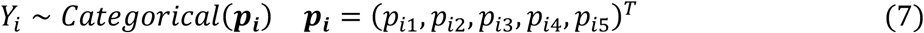

Each parameter is calculated as:

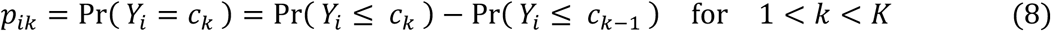

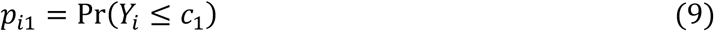

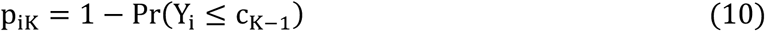

Finally, we compute the cumulative probabilities using ordinal logistic regression:

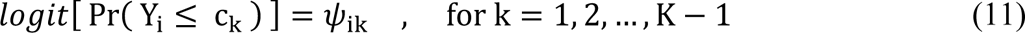

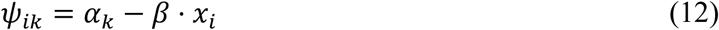

where *x*_*i*_ denotes the library size (total number of reads) for the observation *i*.

This generative model for the observed abundances *Y*_*i*_ is a case of ordinal logistic regression ^32,48^. We use a logit link over the linear predictor ϕ_*ik*_ to estimate cumulative probabilities, i.e., 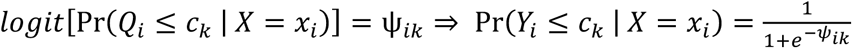. The estimated cumulative probabilities originate the categorical parameters, and the resulting distribution then generates the observed data.

The linear predictor Ψ_*ik*_ has two unknown parameters, the intercepts *α*_*k*_ and the slope β. We have placed weakly-informative priors on both, with no prior preference for any class *c*_*k*_:

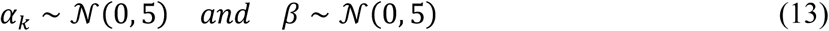

The intercepts are often called cutpoints as they represent the intersections between observable categories on the cumulative logit scale ^49^. Notice we set the same prior for all *K* − 1 cutpoints. The negative-valued slope parameter β seen in equation (12) arises naturally from the PO model derivation with latent continuous variable motivation. It also guarantees intuitive interpretations: positive values indicate a positive effect towards higher categories ^50^.

The ordinal model also allows going beyond conditional (cumulative) class probabilities to estimate conditional expectations, quantiles, and tail probabilities ^32^. This is a major advantage of CPMs over other more commonly used methods such as linear and quantile regression ^30^. We fitted the model using brms and Stan ^33,45^.

### Hierarchical CPM for absolute bacterial abundances

We develop a cumulative logit random effects model to predict bacteria-specific abundances based on observed NGS reads, which is basically a multilevel version of the previous model (7) ^34^. Let *Y*_*ij*_ denote the absolute abundance (in Colony-forming units) for the observation *i*, taxon *j*. Given our serially-diluted samples, we only observe *K* = 4 abundance values such that *Y*_*ij*_ takes values *c*_*k*_ ∈ {*c*_1,_ *c*_2,_ *c*_3_, *c*_4_} = {2 × 10^2^, 2 × 10^3^, 2 × 10^4^, 2 × 10^5^}. We then define the model:

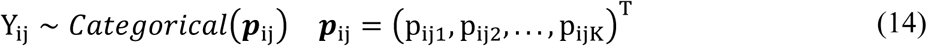

Except for the taxon subscript *j*, the parameters are computed according to equations (8) through (10), and the ordinal regression becomes:

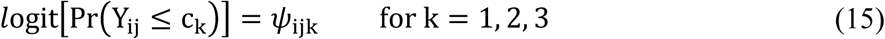

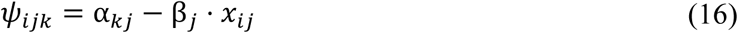

where *x*_*ij*_ is the number of reads in observation *i* from bacteria *j*. Differently from the previous model, here we have only four classes (*K* = 4) and hence *K* − 1 = 3 cutpoints. We allow both intercepts and slopes to vary across bacteria, such that:

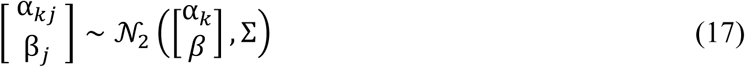

Notice the mean of this two-dimensional Gaussian distribution is the vector of population-level parameters (α_*k*_ *β*)^T^. Thus, the variance-covariance matrix Σ governs how the group-level parameters vary around the population-level counterparts:

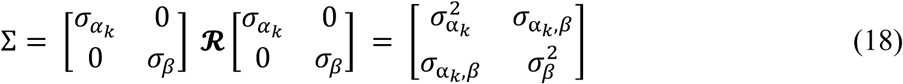

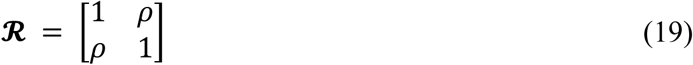

We set the prior distributions for each unknown parameter:

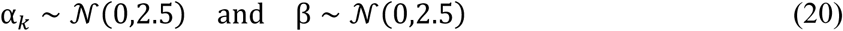

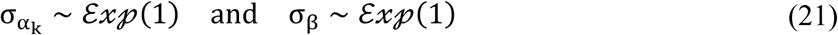

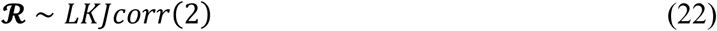

The LKJ prior on the correlation matrix **ℛ** (which describes the correlation ρ between α_*k*_ and β) drives skepticism regarding extreme values near −1 and 1 ^51^. Jointly, the behavior of the prior distributions slightly favors outers categories (*k* ∈ {1, *K*}) in order to improve distinguishability for cases in which there were overlapping average number of reads. The prior choice was driven by model comparison using approximate leave-one-out cross-validation as well as prior and posterior predictive checks ^35,36^.

## Author contributions

G.N.F.C and L.F.V.O performed the bioinformatic analysis. G.N.F.C did the modelling and statistical analysis. A.P.C performed the laboratory experiments. G.N.F.C and A.P.C wrote this manuscript and L.F.V.O performed the revisions.

## Data availability

All sequence data are deposited in NCBI BioProject PRJNA603167.

## Supplementary material

**Supplementary table 1. Sequencing information.** Metrics and specifications about samples and sequencing run data used in this study.

**Supplementary table 2. R packages.**

**Supplementary figure 1.** DNA extraction methods are not a limiting factor in the equivolumetric protocol for bacterial absolute abundances recovery in NGS sequencing.

**Supplementary material 1. Data normalization.**

**Supplementary material 2. Ordinal Regression predicts total microbial load on CFU scale.**

**Supplementary material 3. Ordinal Regression predicts absolute bacterial abundance on CFU scale.**

