## Supplementary Figure 1 for "Equivolumetric protocol generates library sizes proportional to total microbial load in next-generation sequencing"

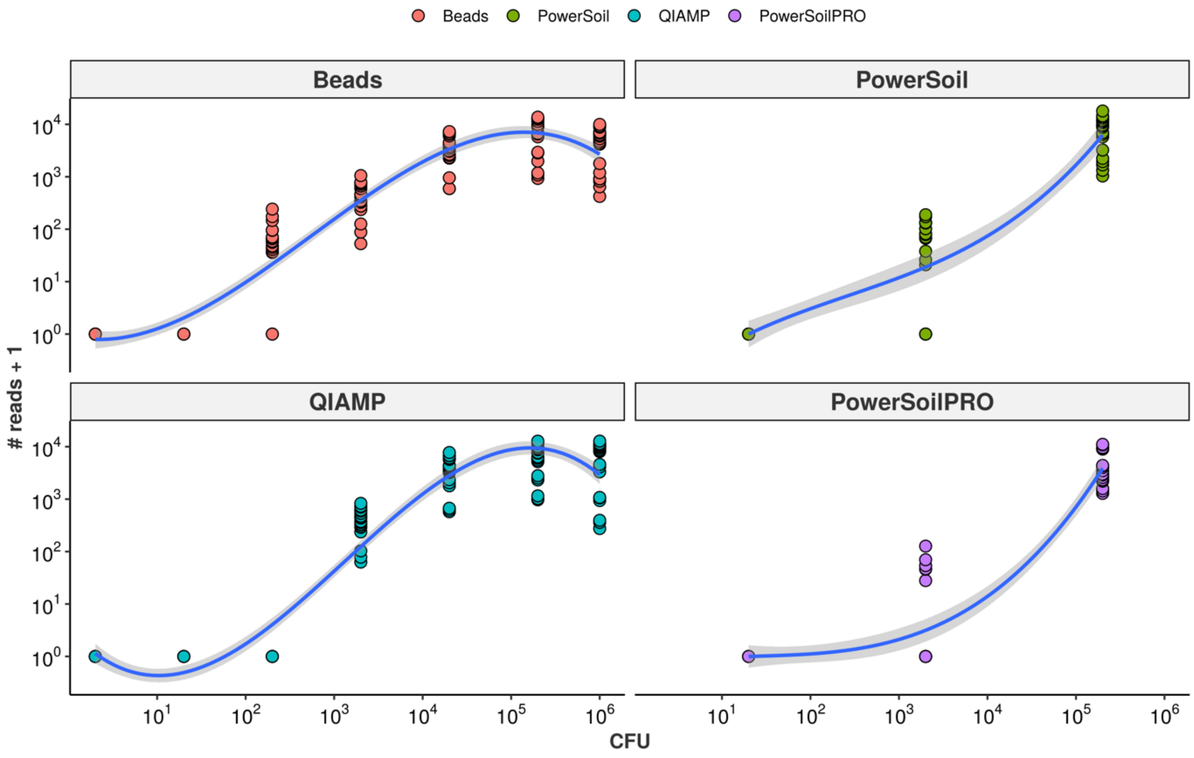


**Supplementary figure 1. DNA extraction methods are not a limiting factor in the equivolumetric protocol for bacterial absolute abundances recovery in NGS sequencing.** Four different DNA extraction methods were evaluated: **Beads** (Magnetic Beads - Agencourt AMPure XP - purification after thermal lysis) (Beckman Coulter, CA, USA); **QIAMP** QIAamp DNA Mini and Blood Mini; **PowerSoil** DNAeasy Power Soil and **PowerSoilPRO** DNAeasy Power Soil PRO (QIAGEN, Germany). For PowerSoil and PowerSoilPRO not all CFU concentrations were evaluated (8.4 CFU to $8.5x10^6$), only three ones representing the lowest, medium and highest values were tested. Overall the results indicated that regardless of the DNA extraction method, the NGS sequencing reads recovers the initial microbial load of the samples when using the equivolumetric protocol.
