## Supplementary Material 1 for "Equivolumetric protocol generates library sizes proportional to total microbial load in next-generation sequencing"

February, 2020

### Supplementary Material 1: Data Normalization

#### Normalization

One of the main challenges regarding the application of NGS technology is the variability of data generated in different sequencing runs, i.e., between-sequencing variation can be significantly driven by batch effects (Robasky, Lewis, and Church 2014). For any given set of samples, different runs commonly yield varying library sizes, even if the runs are performed under equivalent conditions. More importantly, in this case, the number of reads made available *a priori* (expected sample coverage, a measure of relative availability) may vary by significant amounts. One can control such effects, however, while assuming the availability of reads as an upper bound sufficient to exhaust the given libraries - there are more than enough reads available to extract virtually all sample information. Such an assumption is commonly met for samples with low/medium microbe biomass, *e.g.*, hospital microbiome samples with an expected coverage of at least 30000 reads/sample. Otherwise, one must either dilute samples or adjust analysis accordingly. The normalization procedure herein proposed employs a size factor proportional to the expected sample coverage of reads set *before* each sequencing run, representing the relative availability defined for the pool of interest.

##### Size Factor and Scaling

Suppose we have data from multiple sequencing runs with which we want to estimate absolute abundances. In each of the runs, we process a *pool* $p$ of $n$ samples of interest, and the final dataset is a collection of these. Not every pool has the same expected sample coverage. Hence, the normalization proposed is a function of the raw sequenced reads obtained in each run rescaled by a size factor dependent on their relative availability before the sequencing.

$$K^{norm}_{i,j\in D} = \frac{K_{i,j\in D}}{S_{i,j\in D}}$$

Here, $K^{norm}_{i,j\in D}$ denotes the normalized counts for the taxonomy $i$ sample $j \in D$, where $D$ is the set of samples from the $d^{th}$ sequencing run. The size factor is sequencing-specific (i.e., $S_{i,j\in D} = S_d$) and is calculated as follows:

$$S_{i,j\in D} = S_d = \frac{\overline{A_{p^*,d}}}{\max\limits_{d'=1,2,\ldots,m} \left(\overline{A_{p^*,d'}}\right)}$$

where $\overline{A_{p^*,d}}$ is the expected sample coverage in the pool of interest $p^\star$ within sequencing $d$. The lower the relative availability, the smaller the resulting factor and thus greater the normalized values relative to the raw counts. This is important because there are cases in which not every pool has the same number of reads per sample available, which hampers comparability in terms of absolute abundance. Once normalized, divergences across samples from different sequencing runs, but of similar bacterial abundances, are assumed to arise mostly from sequencing efficiency differences - yet of negligible order of magnitude in the scenario of varying biomass. Figure 1 illustrates the normalization effect on experimental data of known abundances from four sequencing runs.

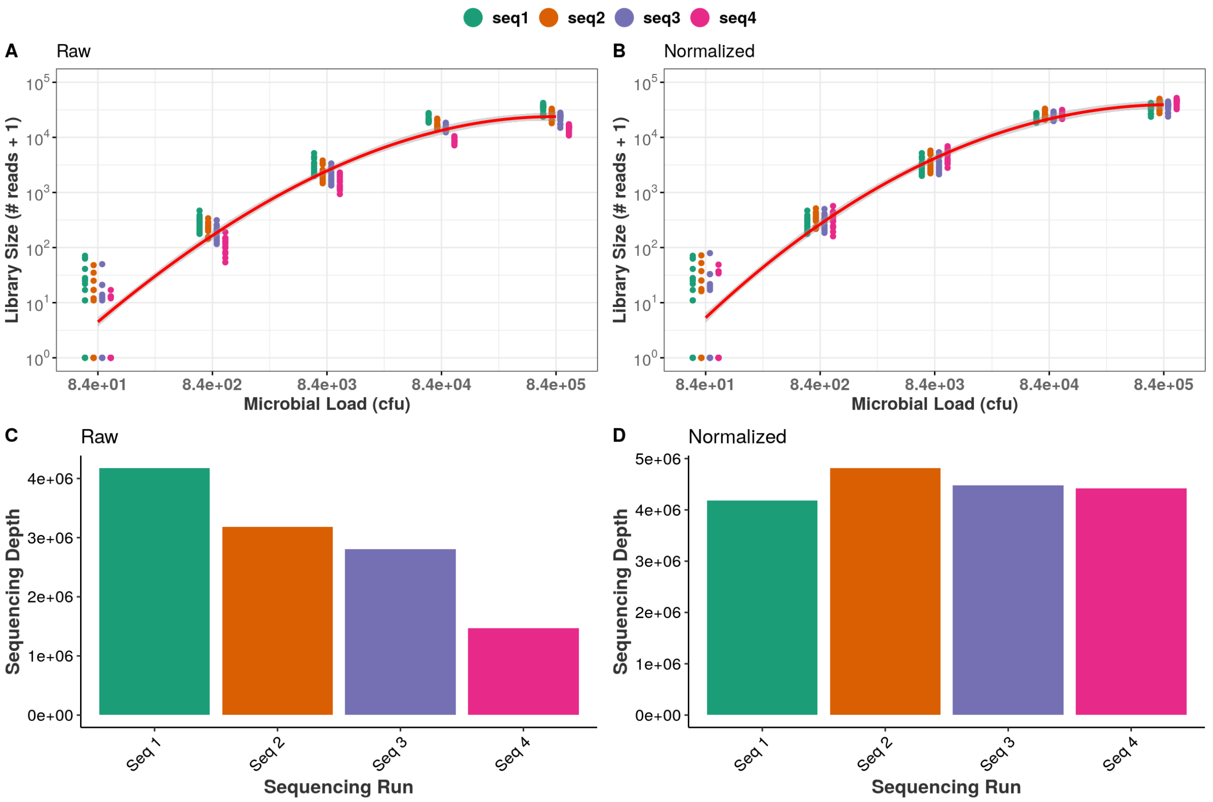


Figure 1. Effect of data normalization.

Notice the third-order polynomial relationship is fit at the log10 scale, even though similar results were observed at the log2 scale (not shown). The former option was preferred mostly because of interpretability: in classical microbiology, log10 is the most common working scale and CFU quantification can be related to orders of magnitude accordingly. The logarithmic transformation also stabilizes variance, addressing issues associated with heteroskedasticity of NGS data.

Beyond such controlled experiments, however, it remains unclear whether the proposed procedure is incapable of fabricating artificial differences. To further validate our approach, we investigated the effect of normalization on a pool of samples sequenced repeatedly in 10-14 different runs from a real hospital environment. Hospital microbiome samples were collected from a Brazillian hospital under the Healthcare-associated Infections Microbiome Project (HAIMP, ethics approval number 32930514.0.0000.0121) (Sereia et al. 2018). Samples (n=180) were collected in April 2015 from Intensive Care Units, Surgical Wards, Internal Medicine Ward, General Surgery Unit, and Emergency room. Bacterial DNA was extracted from samples using a thermal lysis protocol along with posterior AMPure XP magnetic beads (Beckman Coulter, CA, USA). Figure 2 shows the comparison between raw and normalized data.

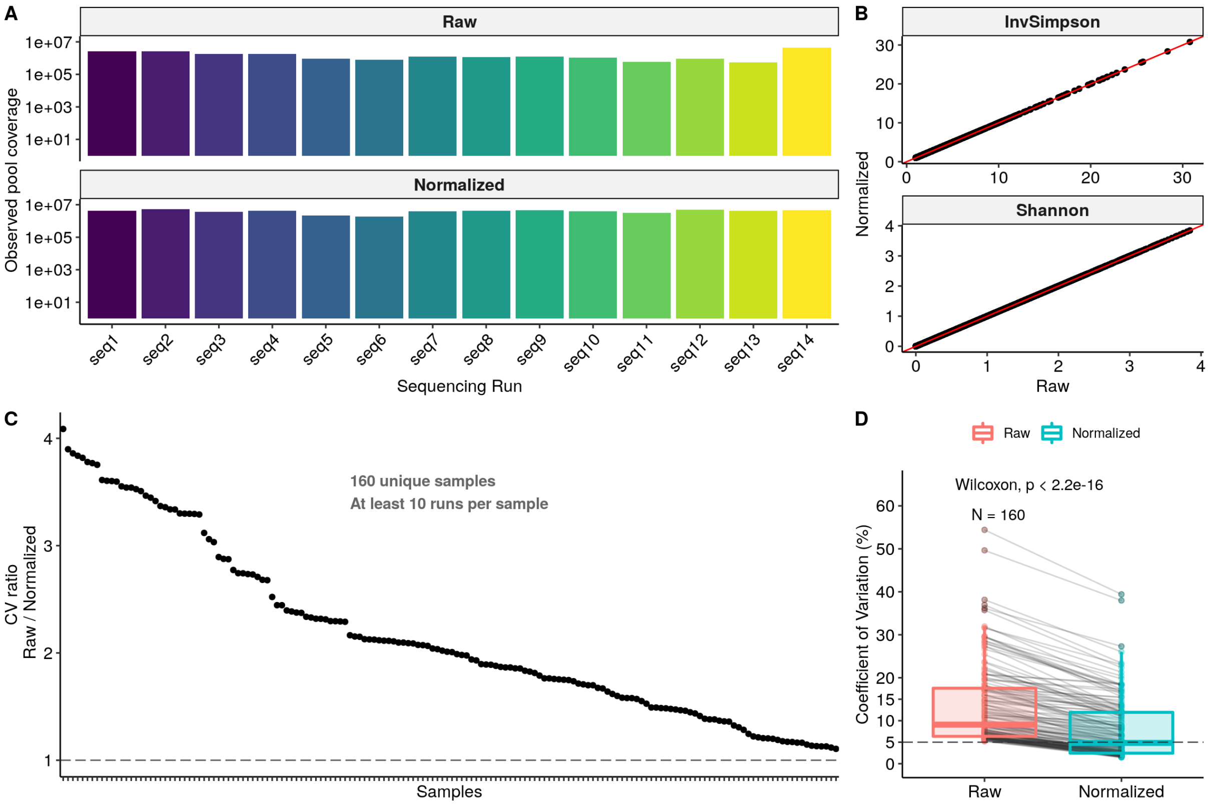


Figure 2. Normalization effects.

In this experiment, the same pool of samples was sequenced in repeated runs (up to 14 times, samples with zero counts were excluded) so that there should be no consistent biological differences across observed pools. Expected sample coverages varied widely (from 60000 to 7891 reads per sample).

The observed pool coverage (Figure 2A) showed little variation even in the raw data, being virtually unaffected after normalization. Figure 2B demonstrates that the procedure does not affect proportion-based metrics such as alpha diversity (red line marks the identity, where $y = x$). Noteworthy, however, is the effect of normalization on sample-wise variability. Using samples sequenced at least 10 times, we computed the coefficient of variation $CV_j = \frac{\hat{\sigma}_j}{\hat{\mu}_j}$ of library sizes for each sample $j$, before and after normalizing. Figures 2C and 2D show that normalized data consistently diminished variability, up to a 4:1 ratio, and brought the median CV down to a 5% level.
