## Supplementary Material 2 for "Equivolumetric protocol generates library sizes proportional to total microbial load in next-generation sequencing"

February, 2020

### Supplementary Material 2: Ordinal Regression predicts total microbial load on CFU scale

This supplementary material addresses model building and validation for total microbial load estimation in terms of Colony-forming Units (CFU). We address the problem in the $log_{10}$ scale as it is commonly used in the realm of classical microbiology. We propose an ordinal regression strategy within the Bayesian framework, although maximum likelihood estimation is also possible.

#### Why ordinal regression?

Colony-forming units are continuous measures. However, in classical microbiology, the ability to quantify microbial abundance has limited resolution: values differ mainly in orders of magnitude, and it is difficult to state replicable differences within the same magnitude range. Hence, decision-making based on such data relies mostly on logarithmic differences, and values such as $2 * 10^3$ and $3 * 10^3$ are often treated as approximately equal. The effect of this "partial discretization" of the outcome space impacts the assumptions of most common regression techniques. Ordinary linear regression is not robust to outliers or high-leverage points, assuming the Gaussian response $Y|X_1$ is a simple shift from $Y|X_2$ (Harrel 2015). Quantile regression is another option but it assumes the distribution of the outcome to be continuous. These challenges hamper the modeling of CFU values and are illustrated with second-order polynomial fits in Figure 1.

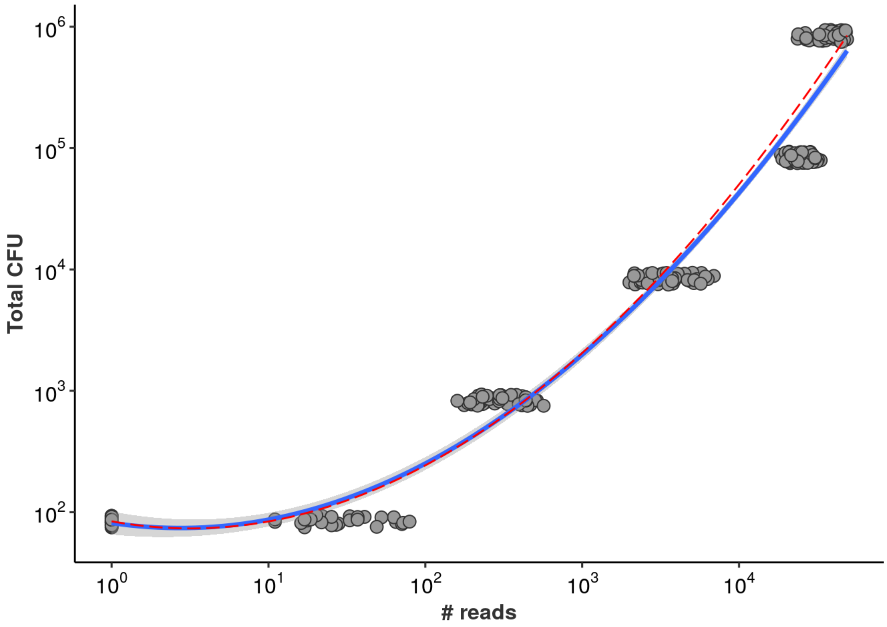


Figure 1. Common regression techniques for estimation of microbial load as a function of library size. OLS regression is shown in blue, and quantile (median) regression in red. A small vertical jittering of the points was used to avoid overplotting.

We show library size on the $log_{10}$ scale (and add a pseudocount of 1 to avoid $log_{10}(0)$). The OLS regression ignores the monotonic, stepwise nature of the relationship between microbial load and library size. Notice the predicted values are not bounded in any way. Higher values of library size will yield estimates well above $10^6$ CFU, which is unrealistic given PCR saturation (see Methods).

The present prediction task, therefore, requires alternatives that are (1) robust to small variations in predictors and response, (2) capable of handling upper- and lower-bounded outcome space, and (3) that respect the monotonic relationship between microbial load and NGS reads in a stepwise fashion. This set of characteristics makes ordinal regression a natural option for predictive modeling.

#### Cumulative Probability Model predicts Total Microbial CFU

Here we describe the proposed *cumulative probability model* (CPM), also called *proportional odds* (PO) model. Formal introductions to ordinal regression are provided in (Bürkner and Vuorre 2019; Liu et al. 2017; Harrel 2015; McElreath 2015). Alternatives to CPM include sequential models and adjacent category models, which we avoid to rely on the interpretability of CPM. Briefly, the model setting is similar to the one in standard (multinomial) logistic regression with $K$ possible outcomes (classes), except that now the log-odds are based on cumulative probabilities. For each class $c_k$ for $k \in \{1, 2, \ldots, K-1\}$:

$$\log \frac{\Pr(Y \leq c_k)}{1 - \Pr(Y \leq c_k)} = \alpha_k - \psi$$

$$\psi = \beta_1 X_1 + \beta_2 X_2 + \cdots + \beta_m X_m$$

The term $\alpha_k$ is the cutpoint associated with the class $c_k$, and the subtracted linear model is left without a standard intercept for identifiability (no $\beta_0$). The subtraction ensures positive coefficients are associated with higher outcome values. When dealing with $K$ total classes, we model explicitly only $K-1$ because the last one is completely determined given a sum-to-one constraint, *i.e.*, $\Pr(Y \leq c_K) = 1$. Note that as we are using only library size as predictor, here $\psi = \beta_1 X_1$. Once the model is fitted, one can recover class probabilities as well as expected values conditional on data:

$$\Pr(Y = c_k \mid X = x) = \Pr(Y \leq c_k \mid X = x) - \Pr(Y \leq c_{k-1} \mid X = x)$$

$$\mathbb{E}[Y|X = x] = \sum_{k=1}^{K} c_k \; Pr(Y = c_k \mid X = x)$$

Calculating conditional expectations may not make sense in many cases (*e.g.* when the response has no continuous interpretation such as $Y \in \{small,\ medium,\ large\}$). Here, on the other hand, expected values may be as interpretable as the classes themselves: when modeling classes of CFU values, we may predict relatively high probabilities to $Y = 1 * 10^1$ and $Y = 1 * 10^2$, for example. In such a case, their weighted average has a clear biological meaning: the expected CFU value is likely to lie between $1 * 10^1$ and $1 * 10^2$. While minimizing the probability of error, the class of highest probability (CHP, the most likely outcome) may suffer in terms of performance measures such as mean squared errors.

### Checking Ordinality Assumption

The main assumption behind the PO model is that the outcome behaves in ordinal fashion with respect to predictors (Harrel 2015). Although Figure 1 suggests monotonicity in the relationship between library size and CFU magnitudes, it is useful to plot stratified averages of the covariate according to levels of the outcome and compare the observations with the model-implied values. We use the function `plot.xmean.ordinaly` from the `rms` package to construct Figure 2.

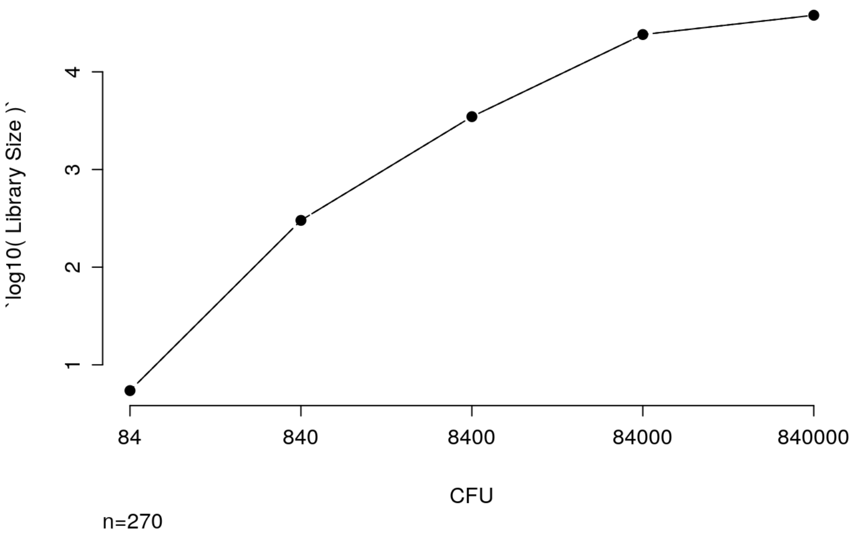


Figure 2. Checking ordinality assumption.

Connected solid dots represent the library size averages ($log_{10}$ scale) stratified by each CFU magnitude level. Assuming that PO holds, the dashed line is the estimated expected value of library size given each value of magnitude (*i.e.*, estimated $\mathbb{E}[\ \mathcal{W} \mid Q = k]$). As the sample means virtually follow the model-implied expectations (almost complete overlap between dashed and solid lines), the graph suggests no major departures from ordinality assumption.

### Model specification

Let $Y_i$ denote the total microbial load (in CFU scale) from the $i^{th}$ sample. Given our serially-diluted samples, we only observe $K = 5$ abundance values such that $Y_i$ takes values $c \in \{c_1, c_2, \ldots, c_5\} = \{0.84 \times 10^2,\ 0.84 \times 10^3, \ldots,\ 0.84 \times 10^6\}$. We then define the model:

$$\begin{gathered}
Y_i \sim \text{Categorical}(\boldsymbol{p_i}) \quad \boldsymbol{p_i} = (p_{i1}, p_{i2}, p_{i3}, p_{i4}, p_{i5})^T \\
p_{ik} = \Pr(\ Y_i = c_k\ ) = \Pr(\ Y_i \leq\ c_k\ ) - \Pr(\ Y_i \leq\ c_{k-1}\ ) \quad \text{for} \quad 1 < k < 5 \\
p_{i1} = \Pr(\ Y_i \leq\ c_1\ ) \\
p_{i5} = 1 - \Pr(\ Y_i \leq\ c_4\ ) \\
\text{logit}[\ \Pr(\ Y_i \leq\ c_k\ )\ ] = \phi_{ik} \quad , \quad \text{for } k = 1, \ldots, 4 \\
\phi_{ik} = \alpha_k - \beta \cdot x_i \\
\alpha_k \sim \mathcal{N}(0, 5) \quad , \quad \beta \sim \mathcal{N}(0, 5)
\end{gathered}$$

where $x_i$ denotes the library size (total number of reads) for the observation $i$. We choose weakly-informative priors for $\alpha_k$ and $\beta$ with no preference for any particular class $c_k$. This generative model for the observed abundances $Y_i$ is a case of ordinal logistic regression (McCullagh, 1985; Harrell, 2015). We use a logit link over the linear predictor $\phi_{ik}$ to estimate cumulative probabilities, i.e., $\text{logit}[\Pr(Q_i \leq c_k \mid X = x_i)] = \psi_{ik} \implies \Pr(Y_i \leq c_k \mid X = x_i) = \frac{1}{1+\exp(-\psi_{ik})}$. The estimated cumulative probabilities originate the categorical parameters, and the resulting distribution then generates the observed data. The linear predictor $\psi_{ik}$ has two unknown parameters (for which we have placed weakly-informative priors): the intercepts $\alpha_k$ and the slope $\beta$. The intercepts are often called cutpoints as they represent the intersections between observable categories on the cumulative logit scale (Agresti 2015). The negative-valued slope arises naturally from the PO model derivation with latent continuous variable motivation. It also guarantees intuitive interpretations: positive values indicate a positive predictor effect towards higher categories (McElreath 2015). The ordinal model also allows going beyond conditional (cumulative) class probabilities to estimate conditional expectations, quantiles, and tail probabilities (Harrel 2015). This is a major advantage of CPMs over other more commonly used methods such as linear and quantile regression (Liu et al. 2017). We fitted the model using brms and Stan (Bürkner 2017; Carpenter et al. 2017).

#### Prior Predictive Check

The weakly-informative priors are on the scale of log-odds and the ordering of cutpoints is enforced by the model. Such settings are constructed in order not to favor any specific magnitude $k$ *a priori* - although class-specific priors are also possible. The fit below is performed ignoring the likelihood - the data information.

```
##  Family: cumulative
##   Links: mu = logit; disc = identity
## Formula: cfu_ranges ~ lib_size
##    Data: df (Number of observations: 270)
## Samples: 4 chains, each with iter = 4000; warmup = 2000; thin = 1;
##          total post-warmup samples = 8000
##
## Population-Level Effects:
##              Estimate Est.Error l-95% CI u-95% CI Rhat Bulk_ESS Tail_ESS
## Intercept[1]    -5.32     15.93   -35.86    26.01 1.00     5396     4683
## Intercept[2]    -1.56     15.94   -32.25    29.92 1.00     5640     5190
## Intercept[3]     1.41     15.88   -29.41    32.53 1.00     5677     5222
## Intercept[4]     5.02     15.89   -25.73    36.37 1.00     5744     5332
## lib_size        -0.03      4.96    -9.58     9.61 1.00     5575     4970
##
## Samples were drawn using sampling(NUTS). For each parameter, Bulk_ESS
## and Tail_ESS are effective sample size measures, and Rhat is the potential
## scale reduction factor on split chains (at convergence, Rhat = 1).
```

The model fit demonstrates the monotonicity of the cutpoints (here as `Intercept[k]` ). Figure 3 shows the actual densities for the parameters (upper panels) as well as the prior predictive distribution behavior (lower panel).

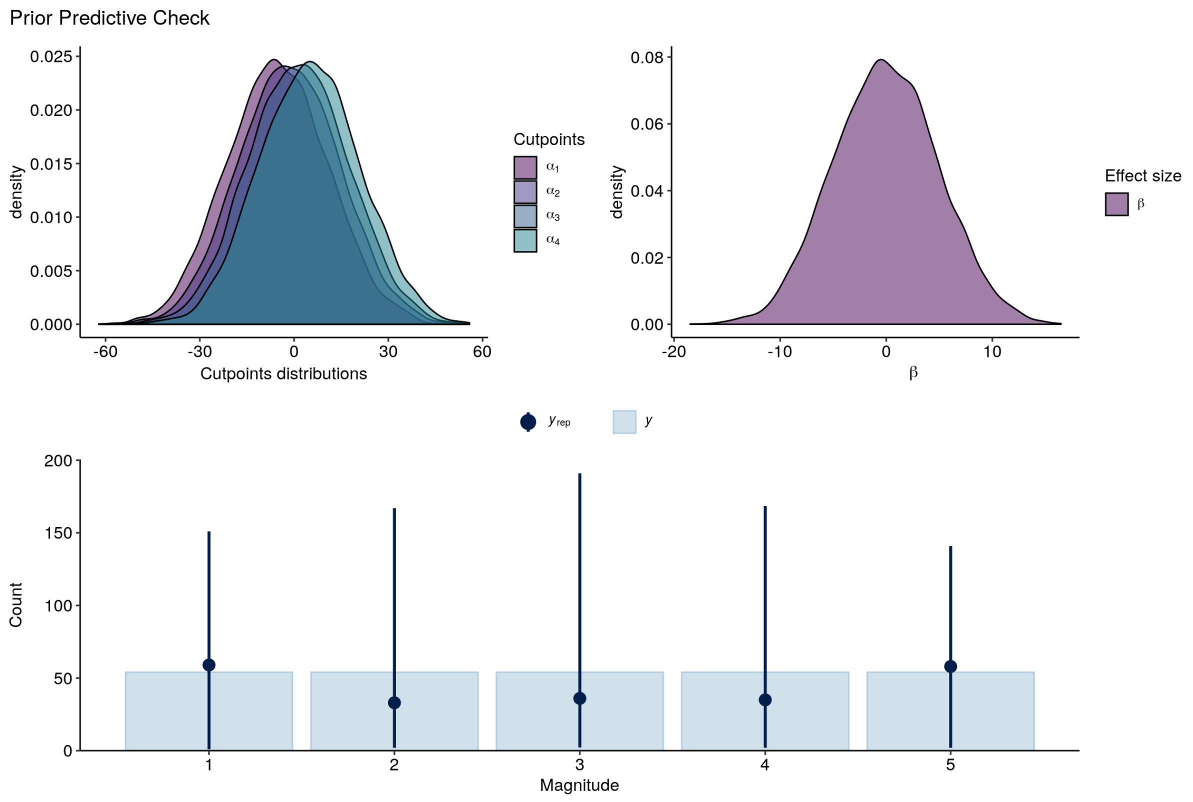


Figure 3. Prior-predictive check indicates incorporation of weak prior information.

The single prior distribution for the cutpoints yields densities that are mostly overlaid, but with slightly increasing locations and high variance. Little information is assumed for the effect of the library size as well. The prior predictive distribution $y_{rep}$ captures the overall structure of the data ($y$) without ruling out most of the outcome space. Notice we have $K-1=4$ cutpoints for $5$ possible values of magnitude.

#### Posterior Predictive Check

Next, we fit the full model (*i.e.* include the likelihood). We observed no signs of lack of convergence in the MCMC algorithm.

```
##  Family: cumulative
##   Links: mu = logit; disc = identity
## Formula: cfu_ranges ~ lib_size
##    Data: df (Number of observations: 270)
## Samples: 4 chains, each with iter = 4000; warmup = 2000; thin = 1;
##          total post-warmup samples = 8000
##
## Population-Level Effects:
##              Estimate Est.Error l-95% CI u-95% CI Rhat Bulk_ESS Tail_ESS
## Intercept[1]    27.34      3.06    21.73    33.66 1.00     2697     4196
## Intercept[2]    40.04      4.22    32.48    48.82 1.00     2415     3242
## Intercept[3]    52.66      5.31    42.99    63.50 1.00     2294     3284
## Intercept[4]    59.35      5.93    48.62    71.51 1.00     2270     2977
## lib_size        13.26      1.33    10.85    15.99 1.00     2270     2975
##
## Samples were drawn using sampling(NUTS). For each parameter, Bulk_ESS
## and Tail_ESS are effective sample size measures, and Rhat is the potential
## scale reduction factor on split chains (at convergence, Rhat = 1).
```

Figure 4 shows the posterior predictive check. The cutpoints and effect of library size are updated to values in the positive real line. The posterior means are well separated. The data (lower panel, $y$) is well captured by the posterior predictive distribution ($y_{rep}$), although there is considerably more variance for the higher classes.

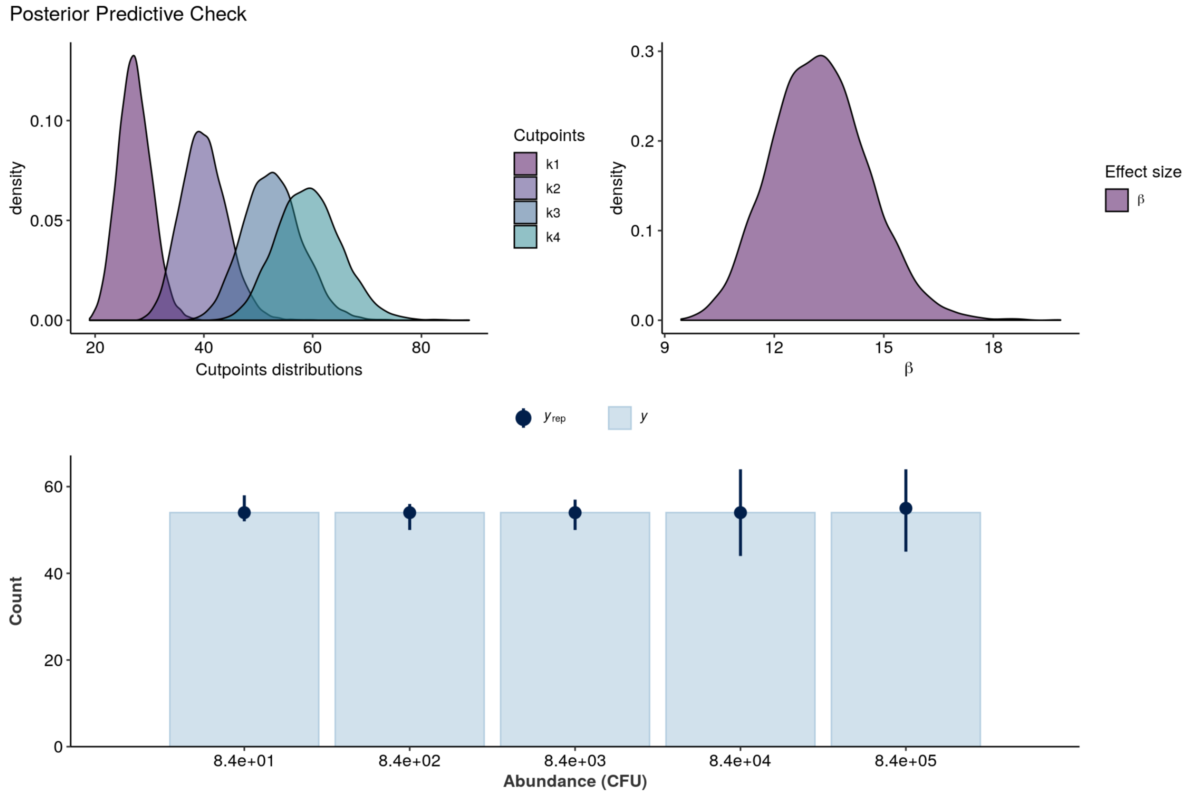


Figure 4. Posterior-predictive check suggests the data-generating process is well captured by the model.

#### CFU Predictions

Given the cumulative probability model structure, we can now recover the CFU magnitude probabilities as well as conditional expectations. We draw fitted values from the (multidimensional) posterior predictive distribution and show the distributions of magnitude probabilities (Figure 5A). These probabilities drive the conditional mean estimates (Figure 5B). Figure 5C shows the predicted values as densities for each outcome value, and Figure 5D replicates the previous OLS regression for comparison.

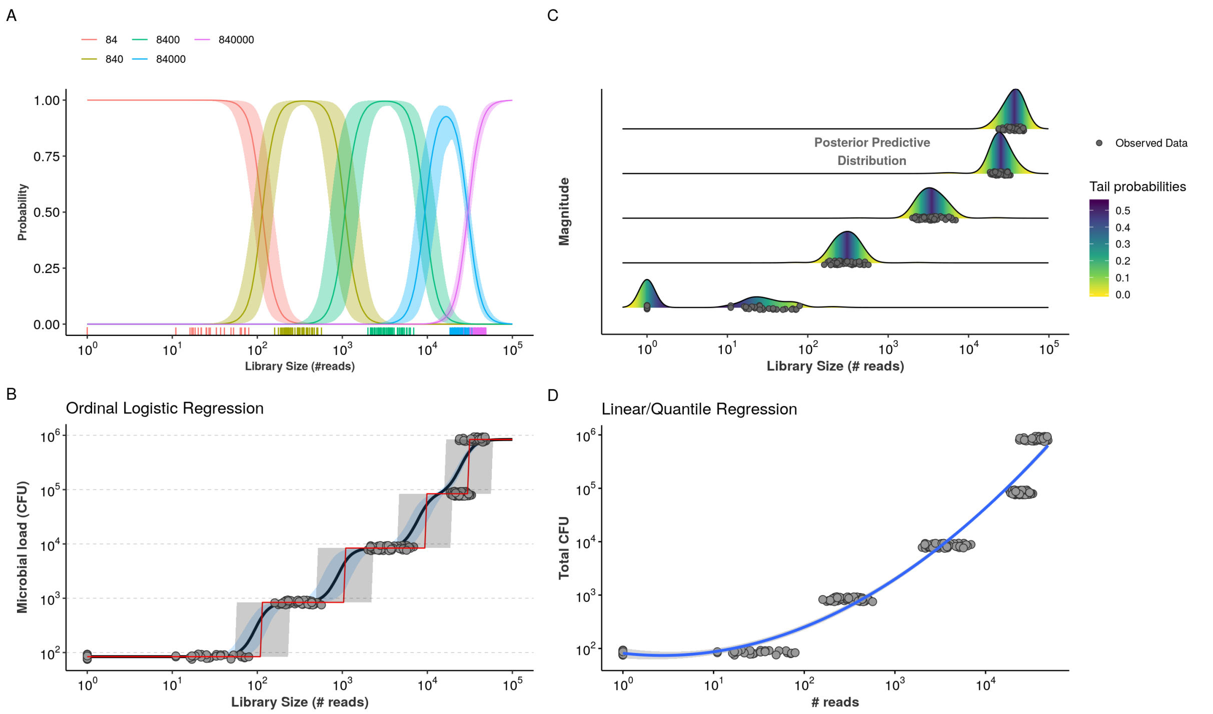


Figure 5. Draws from posterior distribution with respective magnitude probabilities and derived conditional expectations. Figure 1 is reproduced to facilitate comparison.

Note that the estimated mean from the ordinal model follows the stepwise fashion of the data, is bounded within the observed outcome space, and shows similar behavior to standard regression techniques in regions of overlapping read counts. In the CPM, however, the variation is driven by the logistic estimation of magnitude probabilities, rather than direct estimation of expectations. Additionally, the ordinal model properly shows increased uncertainty where there is not much data, suggesting confidence intervals from linear regression may be actually overly confident. Class probabilities are mostly dominant in the non-overlapping regions of the data but still render reasonable posterior predictive distribution.

### Model Validation

We further validate the model using $K$-fold cross-validation to estimate its overall predictive performance, and then use held-out data as a test set.

#### $K$-fold cross-validation

We run $K$-fold cross-validation and the results are shown below. Figure 6A shows the fitted models with respective fold-specific held-out data. Figure 8B shows the predictive performance measures.

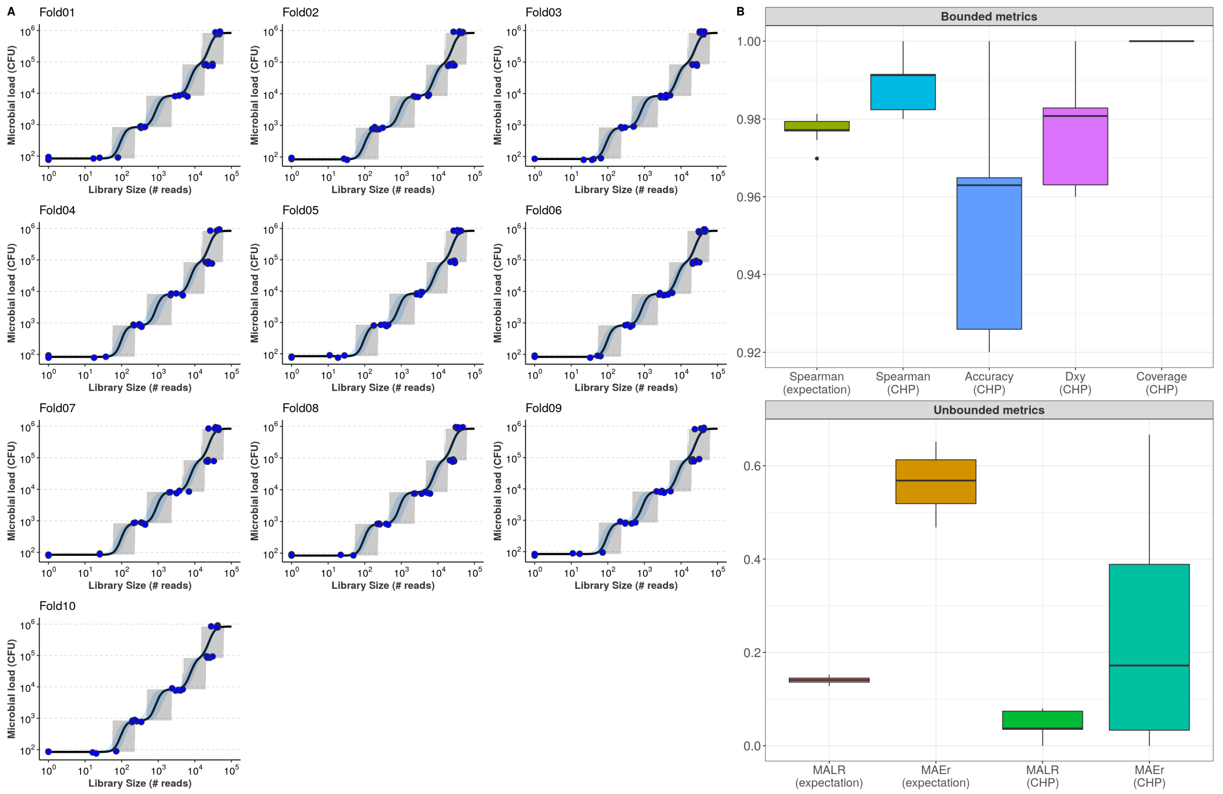


Figure 6. 10-fold cross-validation. (A) CV folds with respective fits and observations. (B) CV performance metrics. ‘CHP’ denotes predictions based on the Class of Highest Probability. Predictions based on expectations are indicated likewise.

For visualization, we have split the assessed metrics into bounded between 0 and 1 and unbounded metrics. Bounded metrics based on CHP included the observed coverage of 95% predictive interval (gray bands in Figure 6A), Somers' Delta (a measure of ordinal association), classification accuracy, and Spearman's rank correlation. The latter was also assessed for expectation-based predictions. In general, these metrics varied well above 0.9. Notably, the predictive intervals showed 100% coverage, which is likely overconfident. Nonetheless, from figure 6A we can see that most intervals spanned only two abundance classes, suggesting errors occur mainly within one order of magnitude from the true values. Ordinal association, as measured by Somers' Delta, was consistently greater than 0.95.

Unbounded metrics relied on modified versions of absolute errors, for both CHP- and expectation-based predictions. MALR denotes Mean Absolute Log-Ratio, a measure that captures how predictions miss observed values in terms of orders of magnitude in the $log_{10}$ scale.

$$\text{MALR} = \frac{1}{n}\sum_{i=1}^{n} \mid log_{10}(\hat{y_i}) - log_{10}(y_i) \mid \quad = \quad \mid log_{10}(\frac{\hat{y_i}}{y_i}) \mid$$

MALR varied during CV below 0.2 for both CHP and expectation. Perhaps more intuitive, we also computed the Mean Absolute Error relative to the true values, capable of measuring absolute errors as proportions of true values.

$$\text{MAEr} = \frac{1}{n}\sum_{i=1}^{n} \frac{|\hat{y_i} - y_i|}{y_i}$$

MAEr tended to be lower for CHP-based predictions compared to expectations, both of each never reaching the value of 0.7.

Notice that a MALR value of 1 corresponds to a ratio between predicted and observed values of one order of magnitude in the $log_{10}$ scale. A MAEr of 1 indicates prediction absolute error as large as the true value, which would still be largely insignificant given the logarithmic scale. Overall, the model generates accurate predictions of total microbial load.

#### Test-set validation

We perform one final model-validation step, which involves predicting new, unseen samples. What we did was holding out beforehand 10% of the dataset to use for such a task. Figure 7 shows the results.

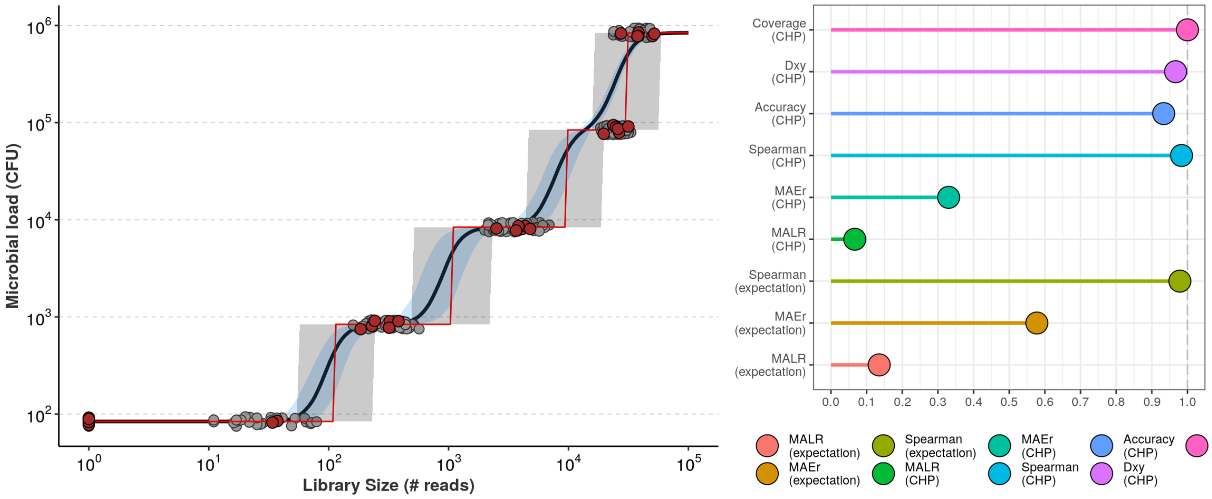


Figure 7, Test-set validation of CPM for total microbial load.

Notice the results from cross-validation are mostly replicated. MAEr values below 0.6, and MALR values below 0.2, indicating predictive errors far below the threshold of one order of magnitude. Predictions show high rank correlations as well as ordinal association with observed values. Classification accuracy seems equally satisfactory.

### Tail probabilities

One of the advantages of the CPM model is the richness of its output. Additionally to class probabilities and conditional expectations, one can retrieve tail probabilities as well (for completeness: one can also calculate conditional quantiles). Once we have $Pr(Y_i = c_k \mid X = x_i)$, we can calculate the probability of having at least $c_k$, conditional on observing $X = x_i$:

$$Pr(Y_i \geq c_k \mid X = x_i) = 1 - Pr(Y_i \leq c_{k-1} \mid X = x_i) = \sum_{j=k}^{c_K} Pr(Y_i = c_j \mid X = x_i)$$

Notice that here we relax the traditional definition of tail probability ($Pr(Y > y)$) to include the class of interest $c_k$.

Figure 8 shows the estimated tail probabilities. Given a library size, each curve shows the probability of having at least $c_k$ CFU for each abundance value.

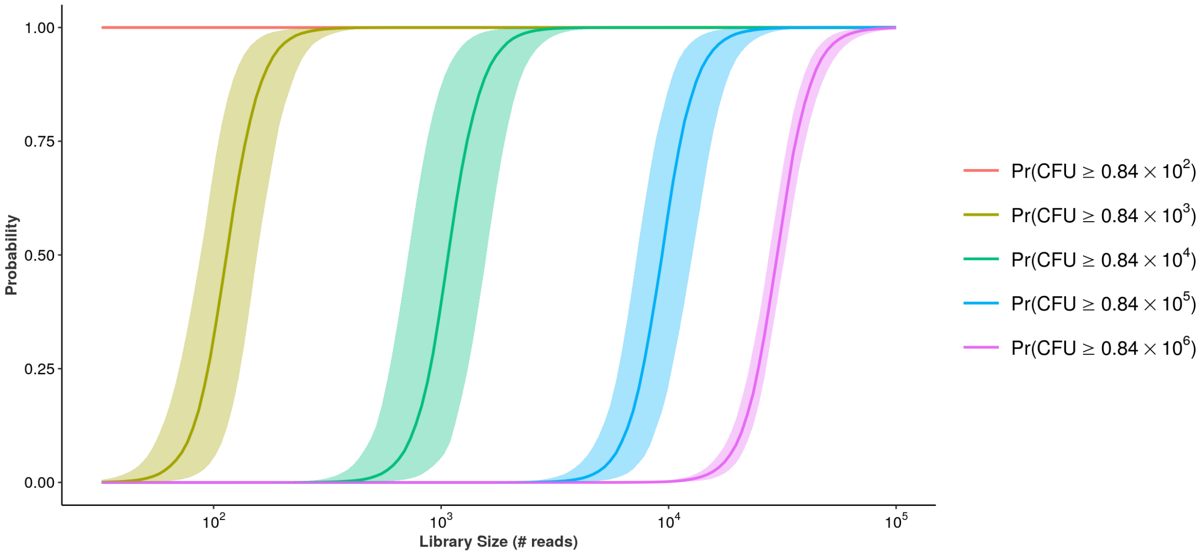


Figure 8. Tail probabilities from CPM model.

The above calculation may have practical applications. For instance, instead of relying on the most likely outcome of derived expectations, it may be enough to know that a sample has a very high probability of having at least $10^4$ CFU of total microbial load. In cases of high estimation uncertainty, one may derive such a lower bound of high probability as:

$$\text{Lower CFU bound} = \max_{1 \leq k \leq K} \{c_k : \Pr(Y_i \geq c_k) \geq \tau\}$$

for some large $\tau$ - say 95% for instance. Analogously, one can describe the samples by the probabilities of having *at most* certain value and hance derive upper bounds of high probabilities:

$$\text{Upper CFU bound} = \min_{1 \leq k \leq K} \{c_k : \Pr(Y_i \leq c_k) \geq \tau\}$$

In the first case, for each observation $Y_i$ you must have $c_k$ or more CFU with high probability (lower bound). In the second case, for each observation $Y_i$ you must have $c_k$ or less CFU with high probability (upper bound).

Finally, it is clear that CPMs can generate a wealth of accurate information useful for total microbial load estimation.
