## Supplementary Material 3 for "Equivolumetric protocol generates library sizes proportional to total microbial load in next-generation sequencing"

February, 2020

### Supplementary Material 3: Ordinal Regression predicts absolute bacterial abundance on CFU scale

This supplementary material addresses model building and validation for the prediction of bacteria absolute abundance in terms of Colony-forming Units (CFU). Similar to the microbial load model, we address the problem in the $log_{10}$ scale as it is commonly used in the realm of classical microbiology. We propose a (Bayesian) mixed-effects cumulative logit model for three main reasons:

- Model the “partially discretized” outcome space as an ordinal variable using cumulative probabilities;
- Accounting for taxon-to-taxon variation in the relationship between number of NGS-derived reads and CFU;
- Take advantage of hierarchical structure and partial pooling.

The first feature addresses peculiarities of the present modeling task which will be further explained below. The taxon-to-taxon variation is a known challenge in the microbiome literature, as different taxa are detected with different efficiencies by NGS technology (McLaren, Willis, and Callahan 2019; Williamson, Hughes, and Willis 2019). Hierarchical structure and partial pooling will be important to allow abundance estimation for previously unseen taxa as well as generalization for out-of-sample data.

For completeness and clarity, some of the material in this section reproduces the text in the Supplementary Material for the microbial load model (Supp. Mat. 2).

#### Why ordinal regression?

Colony-forming units are continuous measures. However, in classical microbiology, the ability to quantify microbial abundance has limited resolution: values differ mainly in orders of magnitude, and it is difficult to state replicable differences within the same magnitude range. Hence, decision-making based on such data relies mostly on logarithmic differences, and values such as $2 * 10^3$ and $3 * 10^3$ are often treated as approximately equal. The effect of this “partial discretization” of the outcome space impacts the assumptions of most common regression techniques. Ordinary linear regression is not robust to outliers or high-leverage points, assuming the Gaussian response $Y|X_1$ is a simple shift from $Y|X_2$ (Harrel 2015). Quantile regression is another option but it assumes the distribution of the outcome to be continuous. These challenges hamper modeling of CFU values and are illustrated with third-order polynomial fits in Figure 1.

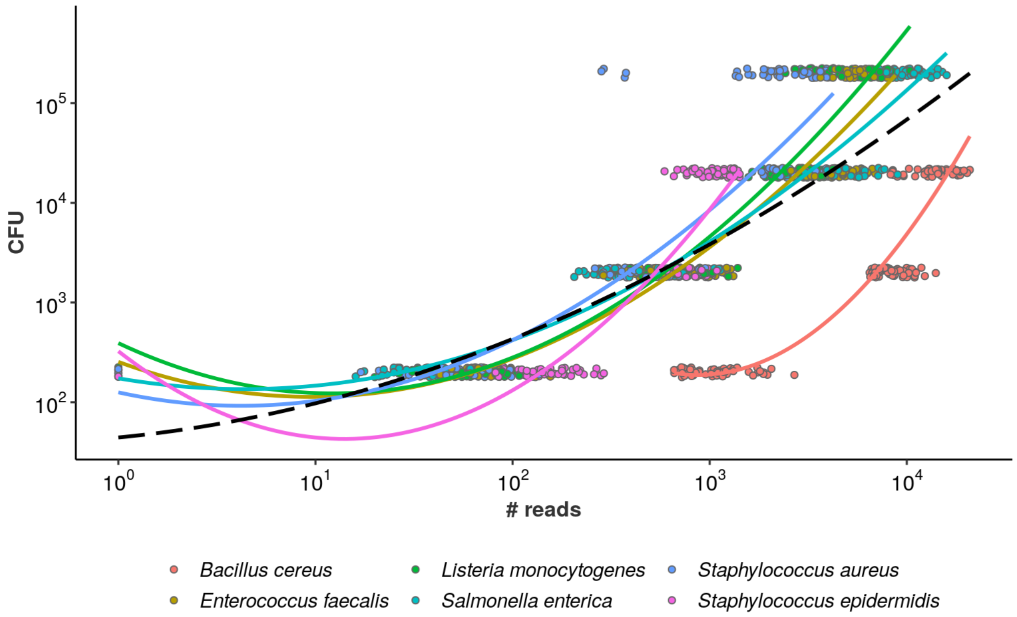


Figure 1. CFU as a function of observed NGS reads (plus pseudocount of 1 to avoid log of zero). Each bacteria is plotted in a different color. Black dashed line indicates overall (“grand mean”) profile. The trend lines were captured with (third-order) polynomial ordinary least squares regression.

Notice how the linear models hardly fit the data. In many cases, the bacteria-specific fits are highly impacted by outliers, especially samples in which the taxon failed to be detected - common situation for low abundances, shown in the plot as $x = 10^0 = 1$ given the addition of pseudocount of 1. Importantly, the predictions are not bounded in any way: higher values of observed reads lead to higher abundances, which is certainly unrealistic given inevitable PCR saturation during laboratory protocol. While the data seems to follow a stepwise fashion (varies mainly across orders of magnitude), the OLS estimator fails to capture such structure. Given this scenario, ordinal models present interesting characteristics:

- **Robust estimation**: we are directly modeling (a function of) the cumulative distribution function (CDF, $Pr(Y \leq y)$) in a semi-parametric manner (W. Liu et al. 2017);
- **Bounded predictions**: as we treat the data as discrete using a multinomial/categorical likelihood for each unique $y_i$ observed, the outcome space is fully determined and all estimations lie within the range of observable data;
- **Ordered modeling of continuous latent variable**: while abundance in CFU is in principle continuous, in practice the observed values differ mainly in orders of magnitude, motivating the derivation of Proportional Odds models (Agresti 2015).
- **Wealth of output information**: from the estimated CDF we can derive conditional values of class probabilities, expectations, quantiles, tail probabilities, etc.

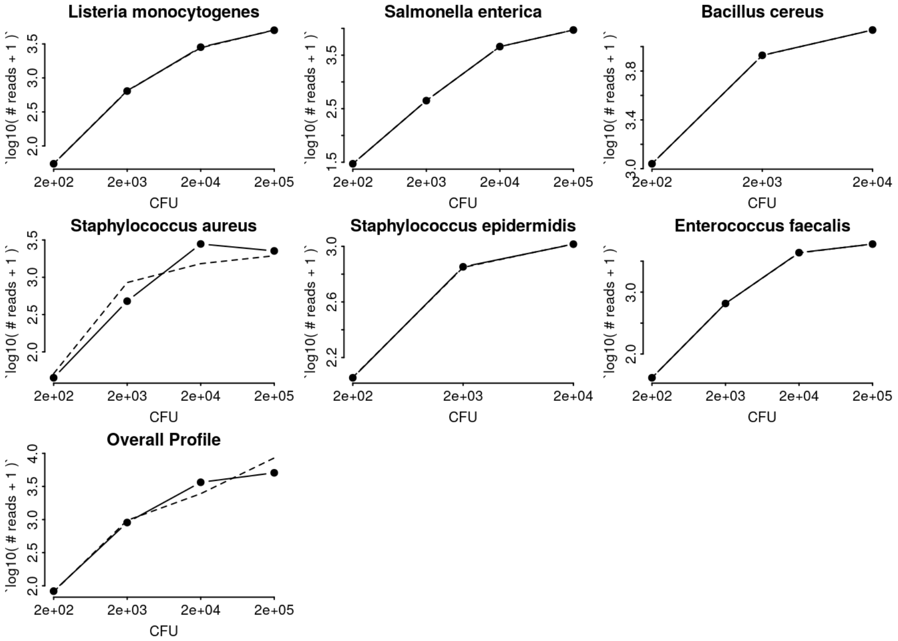


Figure 2. Checking ordinality assumption.

For each bacteria (and overall profile), connected solid dots represent the average NGS reads ($log_{10}(\# \text{ reads} + 1)$) stratified by each CFU abundance level. Assuming that PO holds, the dashed line is the estimated expected value of reads given each value of abundance (*i.e.*, estimated $\mathbb{E}[\ X \mid Y = y]$). As the sample means virtually follow the model-implied expectations, the plots suggest no major departures from ordinality assumptions in most cases, although abundance values $2 * 10^4$ and $2 * 10^5$ might be difficult to distinguish especially for *S. aureus*.

#### Model Specification

We define the cumulative logit random effects model to predict bacteria-specific abundances based on observed NGS reads. Let $Y_{ij}$ denote the absolute abundance (in Colony-forming units) for the observation $i$, taxon $j$. Given our serially-diluted samples, we only observe $K = 4$ abundance values such that $Y_{ij}$ takes values $c \in \{c_1, c_2, c_3, c_4\} = \{2 \times 10^2, \ 2 \times 10^3, \ 2 \times 10^4, \ 2 \times 10^5\}$. We then define the model:

$$Y_{ij} \sim \text{Categorical}(\boldsymbol{p_{ij}}) \quad \boldsymbol{p_{ij}} = (p_{ij1}, p_{ij2}, p_{ij3}, p_{ij4})^T$$
$$p_{ijk} = \Pr(\ Y_{ij} = c_k\ ) = \Pr(\ Y_{ij} \leq\ c_k\ ) - \Pr(\ Y_{ij} \leq\ c_{k-1}\ ) \quad \text{for} \quad 1 < k < 4$$
$$p_{ij1} = \Pr(\ Y_{ij} \leq\ c_1\ )$$
$$p_{ij4} = 1 - \Pr(\ Y_{ij} \leq\ c_3\ )$$
$$logit[\ \Pr(\ Y_{ij} \leq\ c_k\ )\ ] = \phi_{ijk} \quad , \quad \text{for } k = 1, 2, 3$$
$$\phi_{ijk} = \alpha_{kj} - \beta_j \cdot x_{ij}$$

where $x_{ij}$ is the number of reads observed in observation $i$ from bacteria $j$. Differently from the total microbial load model, here we have only four classes ($K = 4$) and hence three cutpoints $\alpha_k$.. More importantly, we allow both intercepts and slopes to vary across bacteria $j$, i.e.:

$$\begin{bmatrix} \alpha_{kj} \\ \beta_j \end{bmatrix} \sim \mathcal{N}_2 \left( \begin{bmatrix} \alpha_k \\ \beta \end{bmatrix}, \Sigma \right)$$

Notice the mean of this two-dimensional normal distribution is the vector of population-level parameters $(\alpha_k \quad \beta)^T$. The variance-covariance matrix $\Sigma$ governs how the group-level parameters vary around the population means:

$$\Sigma = \begin{bmatrix} \sigma_{\alpha_k} & 0 \\ 0 & \sigma_\beta \end{bmatrix} \cdot \boldsymbol{R} \cdot \begin{bmatrix} \sigma_{\alpha_k} & 0 \\ 0 & \sigma_\beta \end{bmatrix} = \begin{bmatrix} \sigma^2_{\alpha_k} & \sigma_{\alpha_k \beta} \\ \sigma_{\alpha_k \beta} & \sigma^2_\beta \end{bmatrix} \quad , \quad \boldsymbol{R} = \begin{bmatrix} 1 & \rho \\ \rho & 1 \end{bmatrix}$$

where $\sigma^2_{\alpha_k}$ is the variance of $\alpha_k$, $\sigma^2_\beta$ is the variance of $\beta$, and $\sigma_{\alpha_k \beta}$ and $\rho$ are their covariance and correlation, respectively. We set the prior distributions for each unknown parameter:

$$\alpha_k \sim \mathcal{N}(0, 2.5) \quad \beta \sim \mathcal{N}(0, 2.5)$$
$$\sigma_{\alpha_k} \sim \mathcal{E}xp(1) \quad \sigma_\beta \sim \mathcal{E}xp(1)$$
$$\boldsymbol{R} \sim \text{LKJcorr}(2)$$

The LKJ prior on the correlation matrix $\boldsymbol{R}$ drives skepticism regarding extreme correlation values near $-1$ and $1$ (McElreath, 2015a). Jointly, the behavior of the prior distributions favors outers categories ($c_k \in \{c_1, c_4\}$) in order to improve distinguishability for cases in which there are overlapping average number of reads.

#### Prior Predictive Check

Our apparently weakly-informative priors actually have an informative joint behavior. We put more weight on outer $c_k$'s in order to improve distinguishability in challenging cases, such as *S. aureus* (see Figure 2). We first fit the model ignoring the likelihood and illustrate such behavior with prior-predictive checks. We then observe the posterior effects of this model in the following section.

```
##  Family: cumulative
##   Links: mu = logit; disc = identity
## Formula: cfu_ranges ~ log10_reads + (1 + log10_reads | bacteria)
##    Data: df %>% mutate(bacteria = factor(bacteria)) (Number of observations: 1188)
## Samples: 4 chains, each with iter = 3000; warmup = 1500; thin = 1;
##          total post-warmup samples = 6000
##
## Group-Level Effects:
## ~bacteria (Number of levels: 6)
##                            Estimate Est.Error l-95% CI u-95% CI Rhat Bulk_ESS
## sd(Intercept)                  1.01      1.00     0.03     3.72 1.00     7505
## sd(log10_reads)                1.01      0.99     0.02     3.67 1.00     7331
## cor(Intercept,log10_reads)    -0.00      0.45    -0.82     0.81 1.00     9177
##                            Tail_ESS
## sd(Intercept)                  3512
## sd(log10_reads)                3195
## cor(Intercept,log10_reads)     3422
##
## Population-Level Effects:
##              Estimate Est.Error l-95% CI u-95% CI Rhat Bulk_ESS Tail_ESS
## Intercept[1]    -2.14      7.81   -17.63    13.66 1.00    10081     4270
## Intercept[2]    -0.04      7.75   -15.42    15.47 1.00    10358     4189
## Intercept[3]     2.09      7.79   -13.02    17.77 1.00    10371     4271
## log10_reads     -0.01      2.56    -5.10     5.16 1.00    10405     4143
##
## Samples were drawn using sampling(NUTS). For each parameter, Bulk_ESS
## and Tail_ESS are effective sample size measures, and Rhat is the potential
## scale reduction factor on split chains (at convergence, Rhat = 1).
```

The model fit, ignoring the likelihood (data), demonstrates the monotonicity of the cutpoints (here as `Intercept[k]` ). Figure 3 shows the actual densities for the parameters (Figure 3A) as well as the prior predictive distribution behavior (Figure 3B).

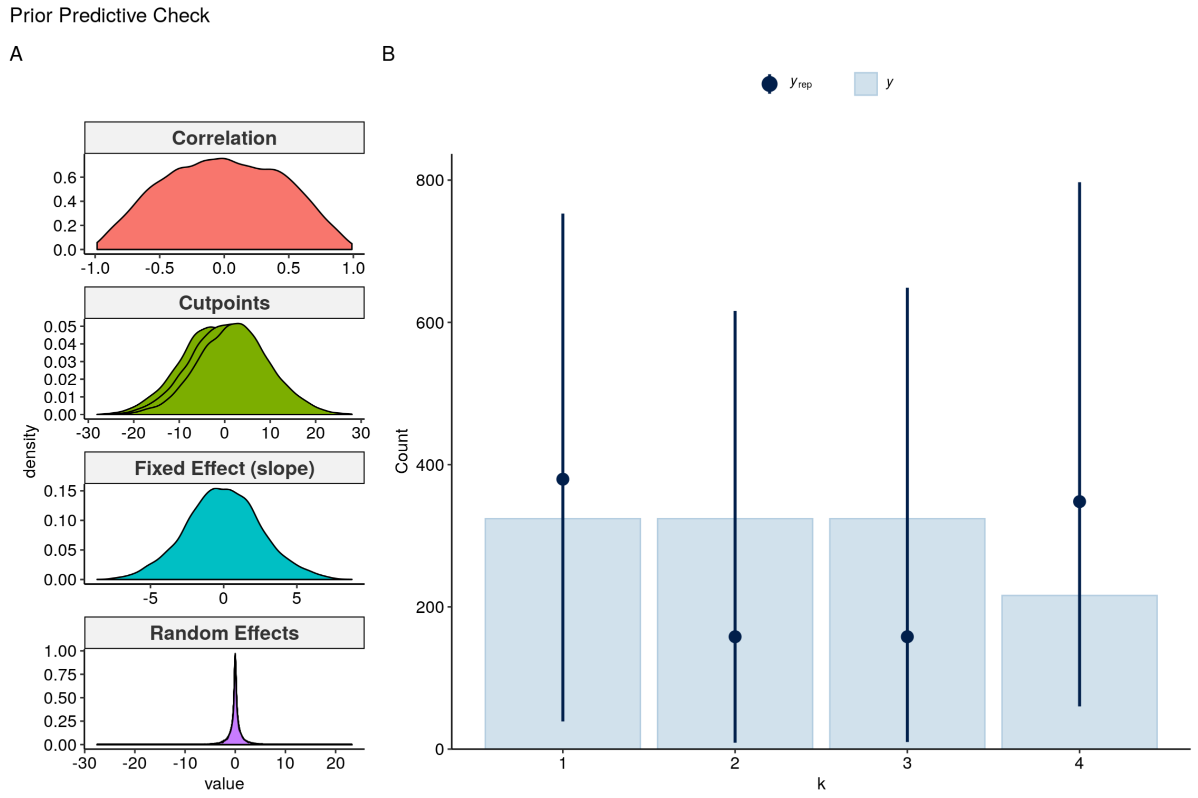


Figure 3. Prior-predictive check.

Notice the priors on the cutpoints (Figure 3B). Although apparently weakly informative ($\mathcal{N}(0, 2.5)$), when applied to the proportional odds model they yield higher (and more variable) probabilities at the outer abundance values ($Y_{ij} \in \{c_1, c_4\}$). While in most cases this information hardly impacts the effect of the likelihood, it will be beneficial to counterbalance cases in which the differentiation between adjacent classes is challenging (*e.g.* higher values for *S. aureus* - see Figure 2). We further explore the consequences of the prior choices in the next section.

### Posterior Predictive Check

Next, we fit the full model (*i.e.* include the likelihood). The overall location of the cutpoints is positive (in logit scale) with relatively small uncertainty. The effect of NGS reads also behaves similarly. Notice intercept and slope are negatively correlated, and the variation in intercepts is considerably higher than that of the slope. The model showed no apparent lack of convergence.

```
##  Family: cumulative 
##   Links: mu = logit; disc = identity 
## Formula: cfu_ranges ~ log10_reads + (1 + log10_reads | bacteria) 
##    Data: df %>% mutate(bacteria = factor(bacteria)) (Number of observations: 1188) 
## Samples: 4 chains, each with iter = 3000; warmup = 1500; thin = 1;
##          total post-warmup samples = 6000
## 
## Group-Level Effects: 
## ~bacteria (Number of levels: 6) 
##                            Estimate Est.Error l-95% CI u-95% CI Rhat Bulk_ESS
## sd(Intercept)                  8.29      1.81     5.30    12.43 1.00     2560
## sd(log10_reads)                2.53      0.67     1.51     4.16 1.00     1864
## cor(Intercept,log10_reads)    -0.61      0.19    -0.88    -0.17 1.00     2467
##                            Tail_ESS
## sd(Intercept)                  3017
## sd(log10_reads)                2619
## cor(Intercept,log10_reads)     3557
## 
## Population-Level Effects: 
##              Estimate Est.Error l-95% CI u-95% CI Rhat Bulk_ESS Tail_ESS
## Intercept[1]    26.28      3.33    19.31    32.31 1.00     1913     2851
## Intercept[2]    35.01      3.47    27.93    41.53 1.00     2092     3082
## Intercept[3]    40.42      3.57    33.26    47.16 1.00     2177     3228
## log10_reads     11.08      1.13     8.74    13.15 1.00     2007     2963
## 
## Samples were drawn using sampling(NUTS). For each parameter, Bulk_ESS
## and Tail_ESS are effective sample size measures, and Rhat is the potential
## scale reduction factor on split chains (at convergence, Rhat = 1).
```

< br/>

Figure 4 shows the posterior distribution of fitted parameters (Figures 4A and 4C) as well as posterior predictive check (Figure 4B).

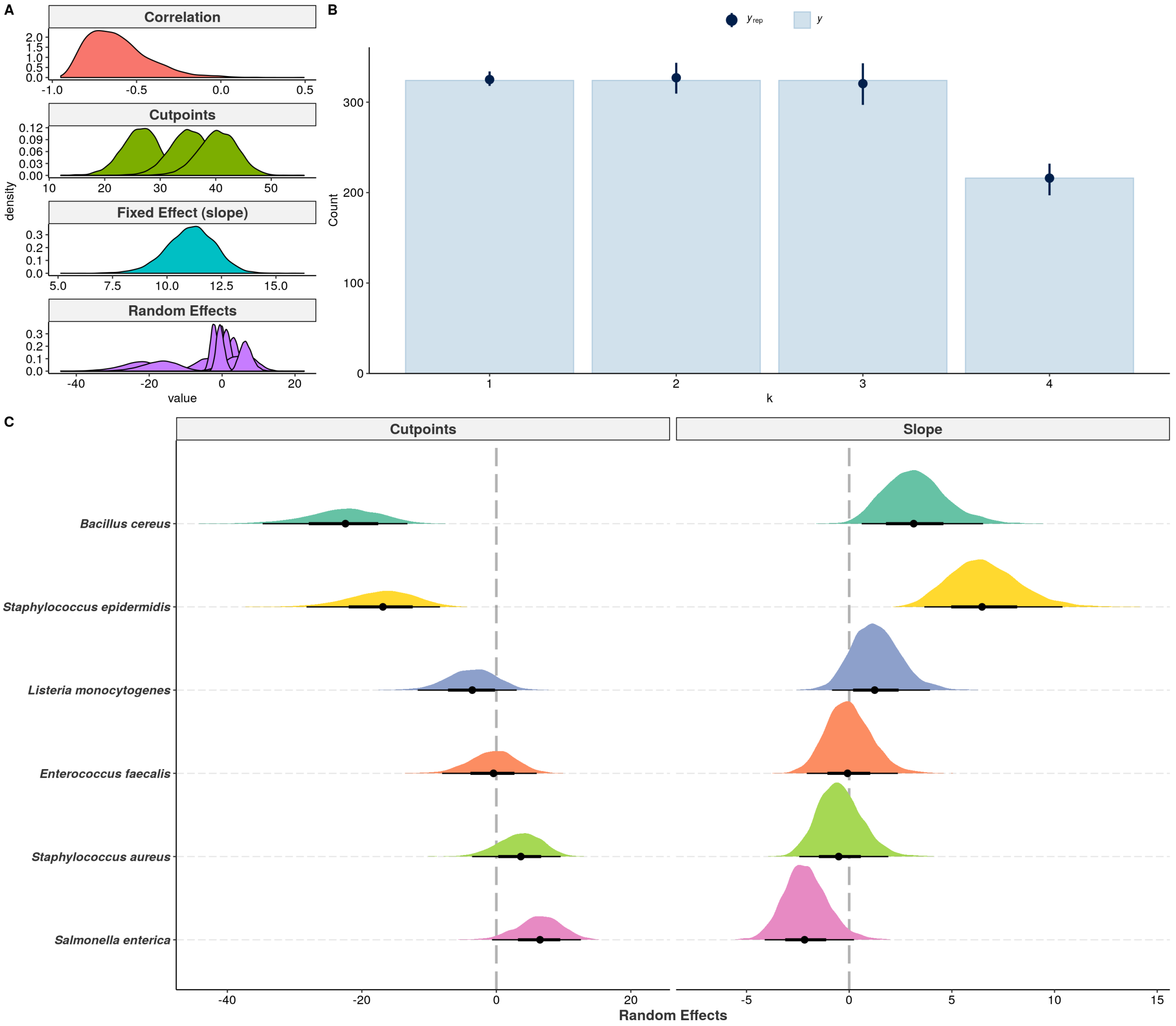


Figure 4. (A) posterior distribution of fixed and random effects. (B) posterior predictive check. (C) posterior distribution of random effects for each bacteria individually. Points show posterior medians, plotted along with 95% and 66% credible intervals.

Overall, the data generating process implied by the model captures the observed data nicely, at least at the population level (Figure 4B). Notice the random effects vary significantly. Under this model, *B. cereus* and *S. epidermidis* seem to be the taxa that most deviate from the overall profile. This is because the posteror distribution of their random effects parameters show the largest absolute values (Figure 4C). Further, we can also perform posterior predictive check in a bacteria-specific fashion, which is shown in Figure 5.

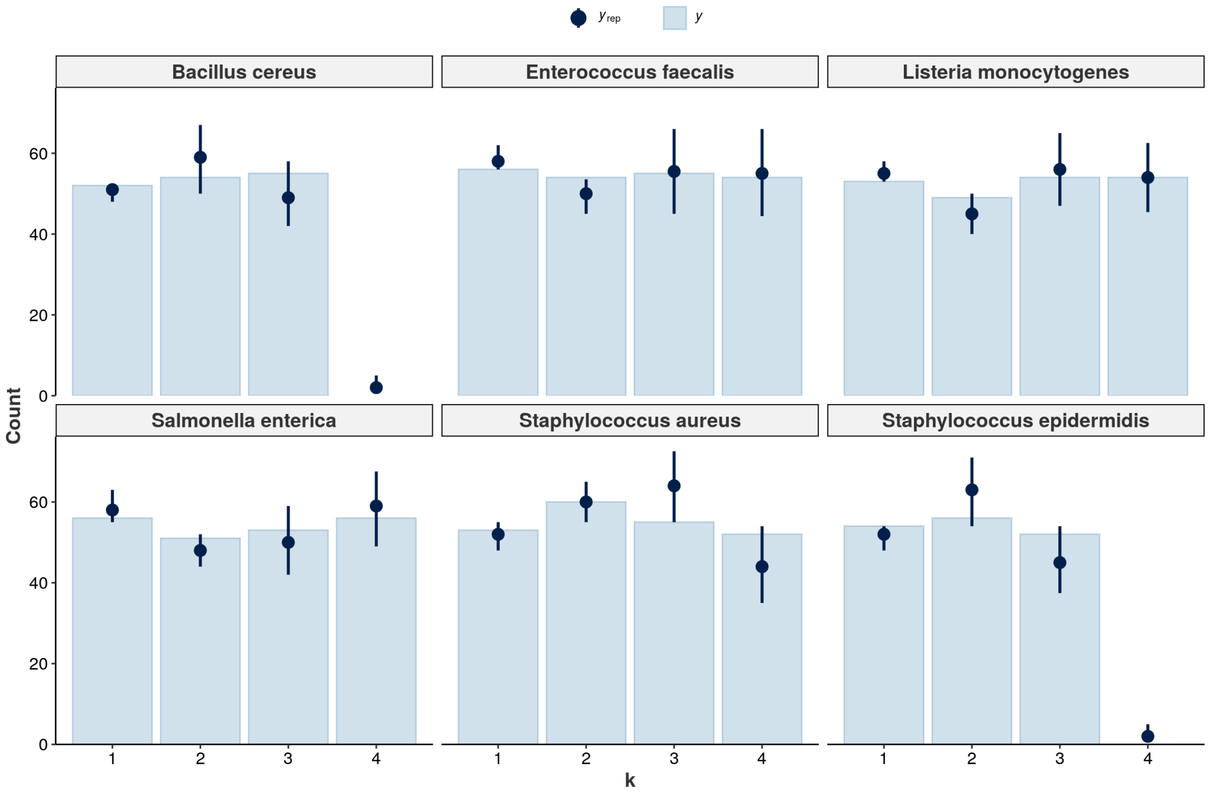


Figure 5. Posterior-predictive check at the bacteria level.

Adaptive shrinkage, characteristic of this type of model, expectedly reduces fit to sample. Yet, the hierarchical model still captures the overall data structure, with variations in some bacteria. Notice the ones that show relatively worse fit are the same that showed paralelism in Figure 2. Especially, $S. aureus$ had shown the least compatible profile, and here yielded the worse posterior fit to the sample - even though the chosen priors mitigate such issue. As there seems to be significant taxon-to-taxon variation, the low number of bacteria available may lead to substantial uncertainty in the estimation of random effects. Nonetheless, while the hierarchical structure is expected to show worse in-sample fit, it improves predictive performance out of sample (McElreath 2015).

### CFU Predictions

Given the cumulative probability model structure, we can now recover the CFU predictions based on class probabilities as well as conditional expectations.

#### Class probabilities

Figure 6 shows the model-implied class probabilities conditional on the number of reads for each observed bacteria.

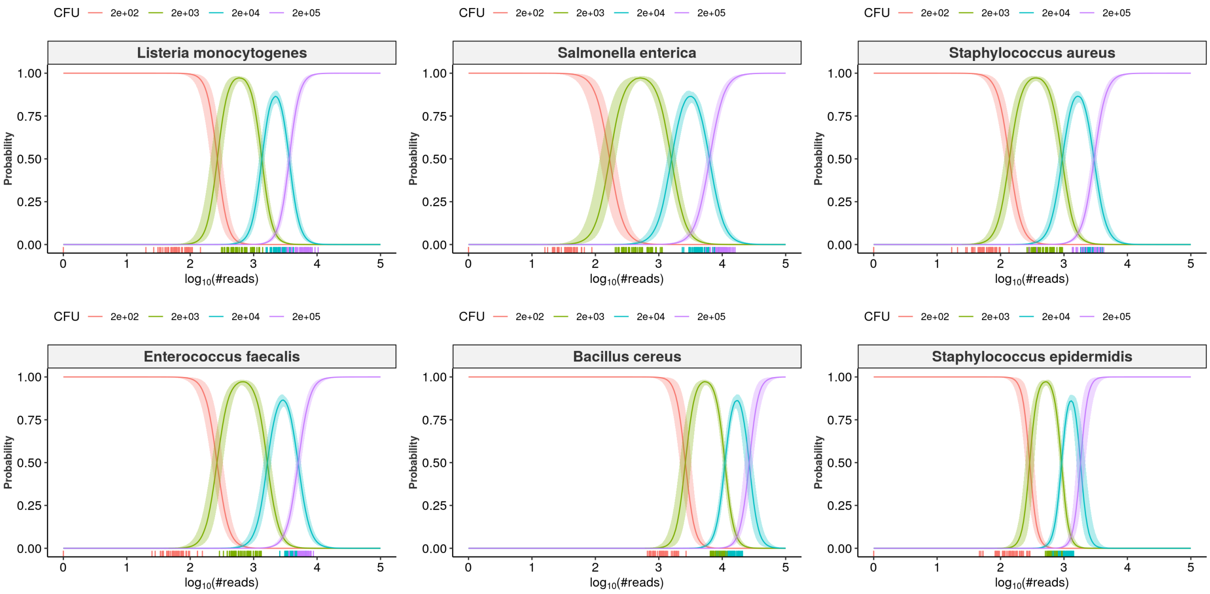


Figure 6. Posterior CFU probabilities as a function of number of NGS reads. Rug plots below each curve show the observed data points.

Notice the profiles differ substantially. Nonetheless, the class probabilities respect the monotonic fashion of the data, *i.e.*, the decrease in probability for a class $c_k$ is accompanied by the increase for class $c_{k+1}$. The rug plots help visualizing the cases in which the model may face difficulty in terms of classification.

#### Expected values and class of highest probability

Next, we compute expected CFU abundances as $\mathbb{E}[Y \mid X = x] = \sum_{k=1}^{K} c_k * Pr(Y = c_k \mid X = x)$ and compare them with the class of highest probability (CHP, most likely outcome for each value $x$). The model-implied CFU predictions are plotted in Figure 7. Black lines show conditional expectations, while red lines indicate the class of highest probability. Respective 95% credible Intervals (CI) are shown in blue and gray.

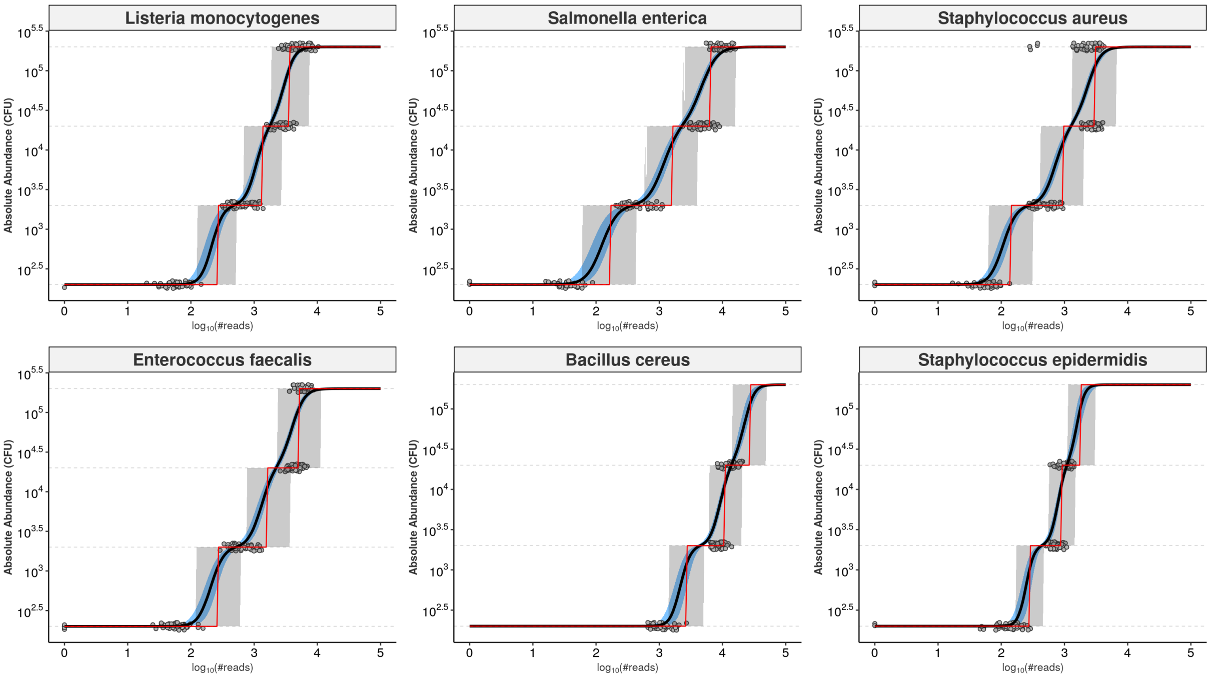


Figure 7. Predictions for Colony-forming units based on NGS reads. Black lines show expected values, while red lines indicate the CHP. Blue bands show the 95% credible interval for the expectations, while gray bands show the 95% predictive interval for the estimated abundance values.

The estimated expectations are smooth functions, while still respecting the monotonic, stepwise fashion of the data. The classes of highest probabilities, in turn, are step functions. Importantly, both are bounded within the outcome space: no predictions fall outside the range of observed data. We replicate parts of Figure 6 and 7 for three bacteria below to stress their relationship with the actual estimated probabilities (Figure 8).

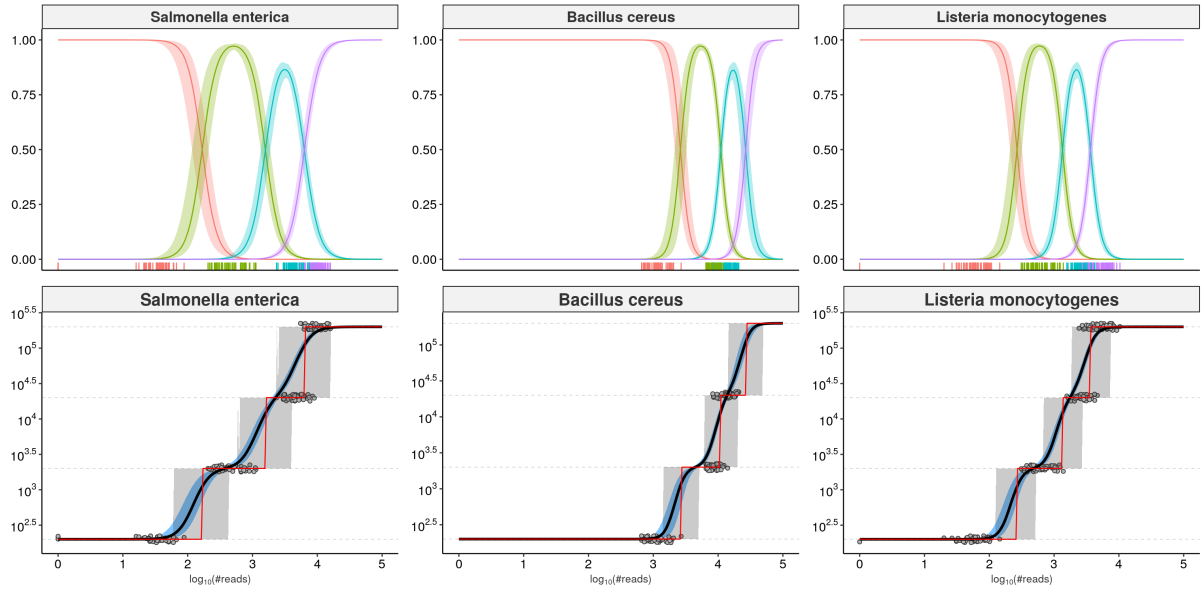


Figure 8. Relationship between estimated probabilities (top) and implied CFU predictions (bottom). Pseudocount of 1 was added to reads values to avoid log of zero.

Notice the strict relationship between the CHP (red lines, lower panels) and the intersections between the class probabilities in the upper panels. Also, the estimated expectations (black lines) vary monotonically as we move away from the probability intersections but remain within the 95% predictive intervals of the abundance values (gray bands).

### Model Validation

We further validate the model using cross-validation and test-set validation with held-out data. First, we perform K -fold cross-validation to estimate overall predictive performance of the model. We then use leave-one-group-out cross-validation to investigate whether the model yields reasonable predictions for previously unseen bacteria.

#### 10-fold cross-validation

We perform 10-fold cross validation and assess the predictive performance. Figure 9 shows the results. For visualization, we have split the assessed metrics into bounded between 0 and 1 and unbounded metrics. Bounded metrics based on CHP included the observed coverage of 95% predictive interval (gray bands in Figure 7), Somers' Delta (a measure of ordinal association), classification accuracy, and Spearman's rank correlation. The latter was also assessed for expectation-based predictions. In general, these metrics had medians varying above 0.9, except for *S. aureus* and *S. epidermidis*. Notably, the predictive intervals showed 100% coverage for almost all bacteria, which is likely overconfident. We also observe median Somers' Delta varying between (slightly below) 0.85 and 0.98.

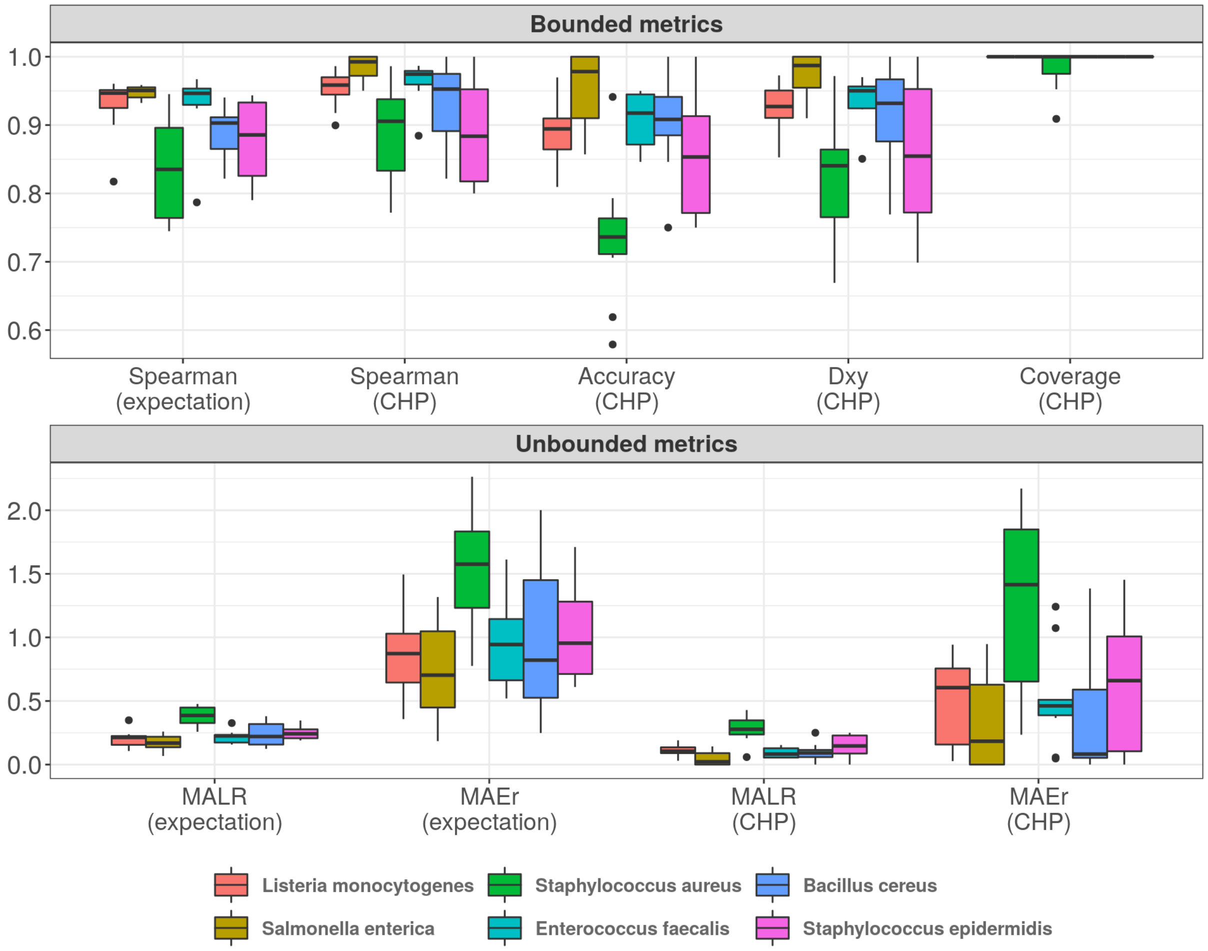


Figure 9. 10-fold cross-validation. 'CHP' denotes predictions based on the Class of Highest Probability. Predictions based on expectations are indicated likewise.

Unbounded metrics relied on modified versions of absolute errors, for both CHP- and expectation-based predictions. MALR denotes Mean Absolute Log-Ratio, a measure that captures how predictions miss observed values in terms of orders of magnitude on the $log_{10}$ scale.

$$\text{MALR} = \frac{1}{n}\sum_{i=1}^{n} \mid log_{10}(\hat{y_i}) - log_{10}(y_i) \mid \quad = \quad \mid log_{10}(\frac{\hat{y_i}}{y_i}) \mid$$

MALR varied consistenly below 0.5 for all bacteria. Perhaps more intuitive, we also computed the Mean Absolute Error relative to the true values, capable of measuring absolute errors as proportions of true values.

$$\text{MAEr} = \frac{1}{n}\sum_{i=1}^{n} \frac{|\hat{y_i} - y_i|}{y_i}$$

Median MAEr values fell below 1 for all bacteria but *S. aureus*, and no MAEr value above 2.5 was observed.

Notice that a MALR value of 1 corresponds to a ratio between predicted and observed values of one order of magnitude in the log10 scale. A MAEr of 1 indicates prediction absolute error as large as the true value, which would still be largely insignificant given the logarithmic scale. Overall, the model seems to generate accurate predictions for all observed bacteria, while *S. aureus* represents the least performing case.

#### Leave-one-group-out Cross-validation

We now explore the predictive performance of our model on estimating previously unseen bacteria through leave-one-group-out cross-validation (LGO-CV). At each iteration, we hold out all the data from one bacteria, train the model with the remaining data, and then predict the held-out partition. The results are shown in Figure 10.

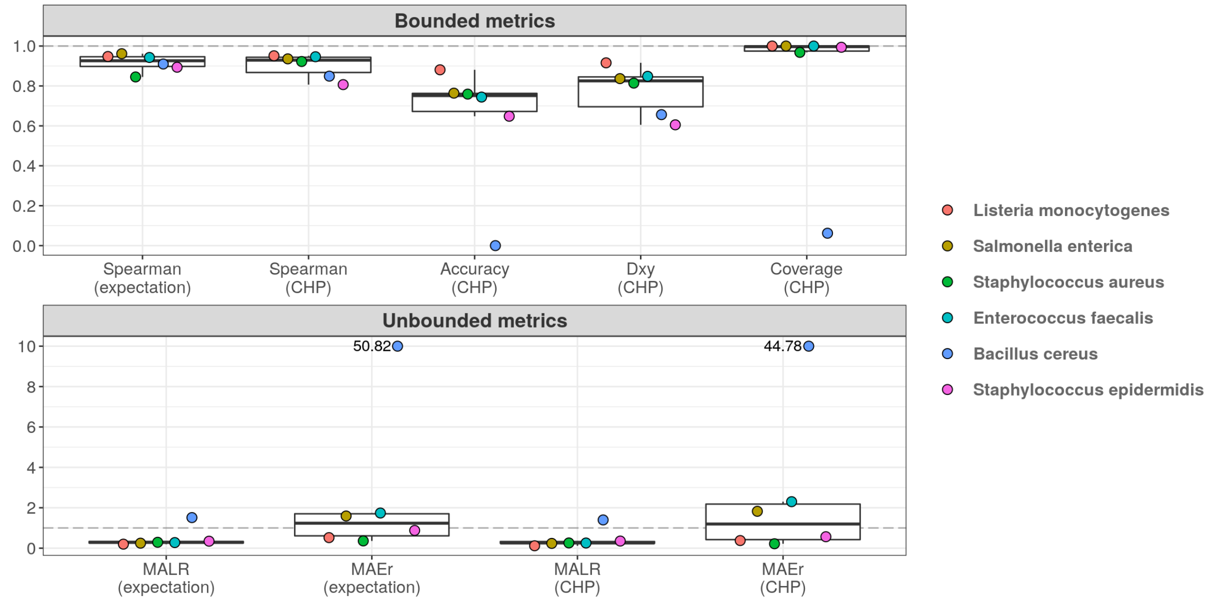


Figure 10. Leave-one-group-out cross validation.

The model completely fails to classify abundance values for *B. cereus*. Yet, for other bacteria, ordinal association and classification accuracy varied between 0.9 and 0.6. Treated as a classification task, the poor predictive performance is likely influenced by high uncertainty associated with the estimation of varying intercepts and slopes using data from only five bacteria at each iteration. This can also explain the high coverage values: 95% predictive intervals were so wide that potentially spanned nearly all outcome space. On the other hand, the taxon associated with the worst out-of-sample performance (*B. cereus*) was also the one with the highest random effects in the original model, *i.e.*, the greatest deviance from the overall, population-level effects - see Figure 4C. While most bacteria showed MAEr between 0.5 and 3, *B. cereus* exceeded the value of 50 (absolute error as large as 50 times the true value). Nonetheless, MALR still varied below the threshold of 1 for all but *B. cereus*, whose associated MALR value almost reached 2 (ratio of two orders of magnitude on the $log_{10}$ scale). It is clear that generalization in high-throughput settings is challenged by specific bacteria, such as *B. cereus*, which deviate greatly from overall profiles. Yet, prediction errors for most bacteria were shown to vary below one order of magnitude.

### Test-set validation

We perform one final model-validation step, which involves predicting new, unseen samples. What we did was holding out beforehand 10% of the dataset to use for such a task. The performance results are shown in Figure 11.

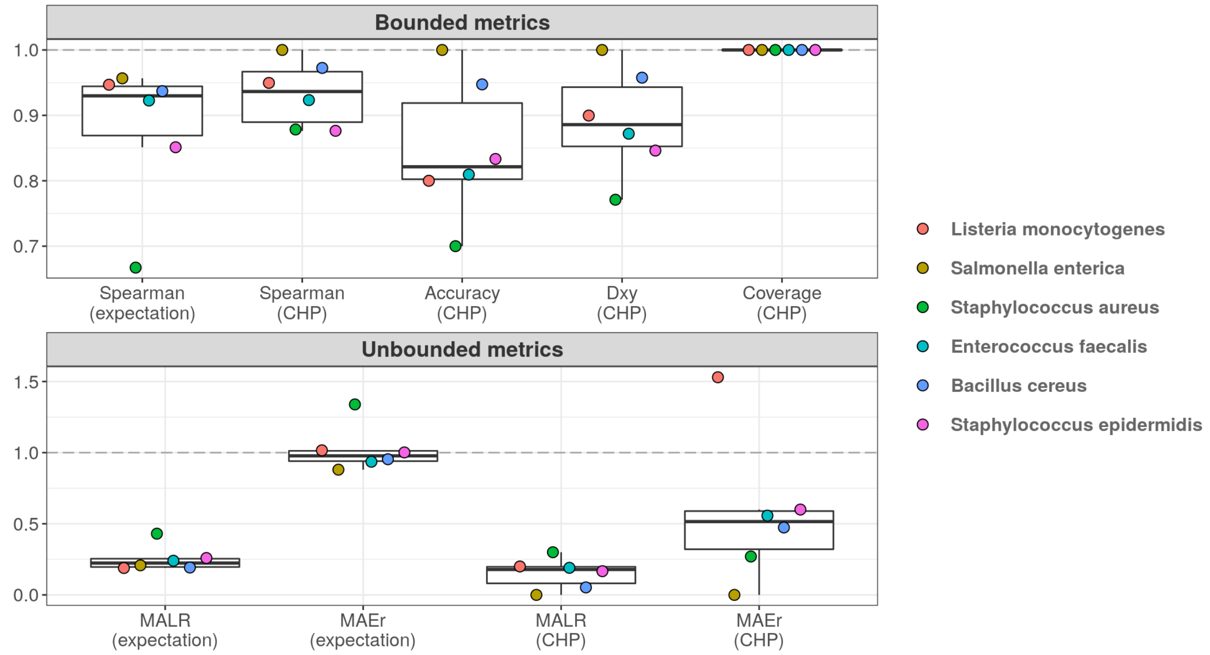


Figure 11. Performance of fitted model on test set.

The highest MAEr value observed was roughly 1.5 (*L. monocytogenes*), while all MALR values were kept below 0.5. *S. aureus* seems to be the taxon with the least preformant profile, agreeing with CV results both in terms of bounded and unbounded metrics. Overall, the results indicate predictive errors remain far below the threshold of one order of mangitude, suggesting that NGS reads can indeed be highly predictive of absolute bacterial abundances.
