## Supplementary Table 1 for "Equivolumetric protocol generates library sizes proportional to total microbial load in next-generation sequencing"

**Supplementary table 1**. Sequencing run information

| Experiment | Seq ID | Date | Kit | Kit Lot | Kit throughput | Reads PF | Clusters PF | Sample Coverage expected | Sample Coverage obtained | PhiX error rate |
|---|---|---|---|---|---|---|---|---|---|---|
| Synthetic fragment | CF47W | 28-Jun-19 | V3-600 | 20339866 20338190 | 25 M | 27,272,423 | 92.48% | 45 | 53.745 | 2.29% |
| HAIMP – Seq1 | AR9G6 | 16-Aug-16 | V2-300 | 20062449 20050998 | 15 M | * | 93.1% | 38.045 | 33.602 | * |
| HAIMP – Seq2 | ARC6K | 2-Sep-16 | V2-300 | 20062449 20050996 | 15 M | * | 93.1% | 29.844 | 40.531 | * |
| HAIMP – Seq3 | ATY5M | 24-Feb-17 | V2-300 | 20081917 20086601 | 15 M | * | * | 28.906 | 29.228 | * |
| HAIMP – Seq4 | AY0WK | 26-May-17 | V2-300 | 20106090 20116131 | 15 M | * | 92.4% | 26.875 | 28.777 | * |
| HAIMP – Seq5 | B47J9 | 20-Jun-17 | V2-300 | 20133531 20140742 | 15 M | * | 94.90% | 26.172 | 21.817 | * |
| HAIMP – Seq6 | AP4Y6 | 16-Sep-16 | V2-300 | 20053902 20047016 | 15 M | * | 90.80% | 25.5 | 17.657 | * |
| HAIMP – Seq7 | AR9J7 | 30-Aug-16 | V2-300 | 20062449 20050996 | 15 M | * | 93.8% | 17.89 | 15.459 | * |
| HAIMP – Seq8 | G19PG | 20-Jun-17 | V2-300 Micro | 20161114 20123715 | 4 M | * | 94.4% | 16.615 | 16.856 | * |
| HAIMP – Seq9 | ARLG5 | 27-Jan-17 | V2-300 | 20067980 20076225 | 15 M | * | 92.0% | 16.485 | 19.428 | * |
| HAIMP – Seq10 | AP6JV | 9-Sep-16 | V2-300 | 20053896 20050996 | 15 M | * | 94.3% | 16.175 | 14.405 | * |
| HAIMP – Seq11 | B449L | 1-Aug-17 | V2-300 | 20133531 20140742 | 15 M | * | 94.1% | 11.745 | 8.475 | * |
| HAIMP – Seq12 | G158L | 22-Feb-17 | V2-300 Micro | 20088731 20126864 | 4 M | * | 94.4% | 11.38 | 12.416 | * |
| HAIMP – Seq13 | AYJPM | 12-Jun-17 | V2-300 | 20123715 20115806 | 15 M | * | 94.4% | 7.891 | 8.886 | * |
| HAIMP – Seq14 | ADC84 | 30-Apr-15 | V2-300 | * | 15 M | * | * | 60 | 61.744 | * |
| ATCC – Seq1 | CGB23 | 26-Jul-19 | V3-600 | 20345776 20351090 | 25 M | 24,440,428 | 90.95% | 45 | 43.527 | 2.61% |
| ATCC – Seq2 | CHW9T | 16-Aug-19 | V3-600 | 20357983 20345731 | 25 M | 23,985,332 | 92.14% | 29.7 | 33.123 | 2.67% |
| ATCC – Seq3 | CJVYY | 23-Aug-19 | V3-600 | 20365653 20360395 | 25 M | 26,025,704 | 88.90% | 28.15 | 29.184 | 2.69% |
| ATCC – Seq4 | CJ3LD | 27-Sep-19 | V2-300 | 20357979 20372379 | 15 M | 13,155,728 | 93.26% | 15 | 15.335 | 2.68% |
| ATCC - Extractions | CKHGJ | 28-Aug-19 | V3-600 | 20369353 20367211 | 25 M | 25,419,260 | 90.76% | 45 | 44.078 | 2.96% |

* missing information

Obs. before 2018 we didn't use PhiX in all the sequencing runs.

Experimental data record improved all over the years, however the results maintained the good yield from 2015 to 2019.

Minimal variations could be caused by the pool quantification and fragment size adjustment.

Error rates and Clusters PF are highly dependent on the sequencing context, giving the majority of sequencing pools are from amplicons, which may lower the sequencing diversity.

Standard sequencing sample coverage for built environments is 45,000, however in these experiments coverages were lowered on purpose to test for diversity an sequencing recovery, as well as the normalization process.

Regardless the sequencing kit used, all runs were performed as single-end 300pb.
