## Supplementary Table 2 for "Equivolumetric protocol generates library sizes proportional to total microbial load in next-generation sequencing"

**Supplementary table 2. References from software packages**

| R Package | Reference |
|---|---|
| brms | Bürkner, P.-C. & Others. brms: An R package for Bayesian multilevel models using Stan. J. Stat. Softw. 80, 1–28 (2017). |
| caret | Kuhn, M. caret: Classification and Regression Training. (2020). |
| DescTools | Signorell, A. DescTools: Tools for Descriptive Statistics. (2020). |
| docstring | Kurkiewicz, D. docstring: Provides Docstring Capabilities to R Functions. (2017). |
| furrr | Vaughan, D. & Dancho, M. furrr: Apply Mapping Functions in Parallel using Futures. (2018). |
| future | Bengtsson, H. future: Unified Parallel and Distributed Processing in R for Everyone. (2020). |
| ggpubr | Kassambara, A. ggpubr: 'ggplot2' Based Publication Ready Plots. (2019). |
| ggrepel | Slowikowski, K. ggrepel: Automatically Position Non-Overlapping Text Labels with 'ggplot2'. (2019). |
| ggridges | Wilke, C. O. ggridges: Ridgeline Plots in 'ggplot2'. (2020). |
| knitr | Xie, Y. knitr: A general-purpose Tool for dynamic report generation in R. R package version 1, (2013). |
| latex2exp | Meschiari, S. latex2exp: Use LaTeX Expressions in Plots. (2015). |
| modelr | Wickham, H. modelr: Modelling Functions that Work with the Pipe. (2019). |
| patchwork | Pedersen, T. L. patchwork: The Composer of Plots. (2019). |
| phyloseq | McMurdie, P. J. & Holmes, S. phyloseq: An R package for reproducible interactive analysis and graphics of microbiome census data. PLoS ONE vol. 8 e61217 (2013). |
| plotly | Sievert, C. plotly for R. (2018). |
| plyr | Wickham, H. The Split-Apply-Combine Strategy for Data Analysis. Journal of Statistical Software vol. 40 1–29 (2011). |
| rafalib | Irizarry, R. A. & Love, M. I. rafalib: Convenience Functions for Routine Data Exploration. (2015). |
| RColorBrewer | Neuwirth, E. RColorBrewer: ColorBrewer Palettes. (2014). |
| Rcpp | Eddelbuettel, D. et al. Rcpp: Seamless R and C++ integration. J. Stat. Softw. 40, 1–18 (2011). |
| readr | Wickham, H., Hester, J. & Francois, R. readr: Read Rectangular Text Data. (2018). |
| rlang | Henry, L. & Wickham, H. rlang: Functions for Base Types and Core R and 'Tidyverse' Features. (2020). |
| rms | Harrell, F. E., Jr. rms: Regression Modeling Strategies. (2019). |
| scales | Wickham, H. & Seidel, D. scales: Scale Functions for Visualization. (2019). |
| splines | R Core Team. R: A Language and Environment for Statistical Computing. (2019). |
| tidybayes | Kay, M. tidybayes: Tidy Data and Geoms for Bayesian Models. (2020) doi:10.5281/zenodo.1308151. |
| tidyverse | Wickham, H. et al. Welcome to the tidyverse. Journal of Open Source Software vol. 4 1686 (2019). |
